# The impact of kit, environment and sampling contamination on the observed microbiome of bovine milk

**DOI:** 10.1101/2023.11.07.566052

**Authors:** C. J. Dean, Y. Deng, T. C. Wehri, F. Pena-Mosca, T. Ray, B.A. Crooker, S. M. Godden, L. S. Caixeta, N.R. Noyes

**Affiliations:** Department of Veterinary Population Medicine, University of Minnesota, St. Paul, 55108; Department of Animal Science, University of Minnesota, St. Paul, 55108

## Abstract

Contaminants can easily outnumber bacteria that originate within the milk itself, milk microbiome research currently suffers from a critical knowledge gap; namely, does non-mastitis bovine milk contain a native microbiome? In this study, we sampled external and internal mammary epithelium, stripped and cisternal milk, used numerous negative controls to identify potential sources of microbial contamination. Two algorithms were used to mathematically remove this contamination and to track potential movement of microbes among our samples. Our results suggest that majority (i.e., >75%) of the sequence data generated from bovine milk and mammary epithelium samples represents contaminating DNA. The contaminants in milk samples were primarily sourced from the DNA extraction kits and the internal and external skin of the teat, while the teat canal and apex samples were mainly contaminated during the sampling process. After decontamination, the milk microbiome displayed a more dispersed, less diverse and compositionally distinct bacterial profile compared with the teat skin samples. Similar microbial compositions were observed between cisternal and stripped milk samples, as well as between teat apex and canal samples. *Staphylococcus* and *Acinetobacter* were the predominant genera detected in the sequences of milk samples, and bacterial culture showed growth of *Staphylococcus* and *Corynebacterium* spp. in 50% (7/14) of stripped milk samples and growth of *Staphylococcus* spp. in 7% (1/14) of cisternal milk samples. Our study suggests that microbiome data generated from milk samples obtained from clinically healthy bovine udders may be heavily biased by contaminants that enter the sample during the sample collection and processing workflows.

**Importance:** Obtaining a non-contaminated sample of bovine milk is challenging due to the nature of the sampling environment and the route by which milk is typically extracted from the mammary gland. Furthermore, the very low bacterial biomass of bovine milk exacerbates the impacts of contaminant sequences in downstream analyses, which can lead to severe biases. Our finding showed that bovine milk contains very low bacterial biomass, and each contamination event (including sampling procedure and DNA extraction process) introduces bacteria and/or DNA fragments that easily outnumber the native bacterial cells. This finding has important implications for our ability to draw robust conclusions from milk microbiome data, especially if the data have not been subjected to rigorous decontamination procedures. Based on these findings, we strongly urge researchers to include numerous negative controls into their sampling and sample processing workflows; and to utilize several complementary methods for identifying potential contaminants within the resulting sequence data. These measures will improve the accuracy, reliability, reproducibility, and interpretability of milk microbiome data and research.

## Introduction

Animals are hosts to a variety of microbial communities, many of which are site-specific, and their structure and function are often unique to a specific anatomical location within the host [1]. These site-specific microbiomes can play important roles in the host’s physiology and health through host-microbe and microbe-microbe interactions [2,3]. In dairy cows, the mammary gland is a critical organ due to its central role in milk production [4,5]. The secretory portion of the bovine mammary gland is anatomically complex, composed of four separate compartments (quarters) that contain secretory epithelial cells (i.e., lactogenic cells) capable of synthesizing, storing, and secreting milk components into individual alveoli from which the milk travels through ducts to the udder cistern. Milk produced from a healthy bovine mammary gland was historically believed to be sterile, particularly at its origin high up within individual mammary alveoli [6,7]. However, even in healthy mammary glands, results from culture-independent molecular approaches have cast doubt on the sterility of *in situ* milk because these results show a diverse microbial community profile obtained from 16S rRNA-based workflows [6,8,9]. The 16S rRNA approach uses sequence variability in certain regions of the 16S rRNA gene to taxonomically classify DNA fragments extracted from a raw sample such as bovine milk [10]. Using this approach, previous studies have reported associations between the bovine milk microbiome and immune responses [11], development of mastitis [4], quality and safety of dairy products [12], and the gut microbiota of both calves and consumers [5,13]. Taken together, this body of literature suggests that the milk microbiome is a crucial component of cow health and production as well as milk safety and quality.

Milk is known to be a low-biomass sample matrix, dominated by non-cellular components such as proteins and fat [14]. Additionally, the cellular components of milk tend to be dominated by host (i.e., *Bos taurus*) cells as opposed to microbial cells [15,16]. Low-biomass samples are especially challenging for microbiome studies because cells and/or DNA introduced during sample collection and processing can easily outnumber endogenous cells and/or DNA. The challenges of low-biomass sample matrices for microbiome studies have been well-documented [17–21]. However, even amongst low-biomass sample matrices, milk has long been noteworthy for the difficulty of obtaining a “clean” sample. Indeed, contamination rates are usually >10% for bovine milk samples subjected to bacterial culture [22]. The reason for this high rate of bacterial contamination is the process by which milk is typically sampled, namely by collecting so-called “stripped milk”. In this sampling procedure, the external skin of a teat is first thoroughly cleaned and disinfected. Then, the sampler manually expresses and discards several streams of milk from the teat. It is thought that this initial milk stream is more likely to contain contaminating bacteria due to the milk flowing through the lower compartments of the udder, i.e., the teat canal, which is more easily accessed by bacteria that can move in a retrograde fashion from the teat apex up into the canal and, if not thwarted by the immune system, from there into the udder cistern, ducts and mammary tissue. Thus, after discarding the initial milk streams, “aseptic” milk is then collected via continued manual expression from the teat [23]. However, even this process cannot eliminate potential contamination of the stripped milk from bacteria and/or DNA located on the external teat epithelium and/or the epithelium that lines the internal teat canal [24,25]. Teat wall puncture is often recommended as the most reliable method of collecting a clean, uncontaminated milk sample [26].

Additionally, even milk not contaminated with bacteria associated with external epithelial can still be contaminated at any of the numerous other downstream harvest and processing procedures, including from exposure to ambient air in the room where samples are being collected. This is a particular concern for most bovine milk microbiome studies, which typically collect samples in large animal handling facilities such as milking parlors or research barns. These environments often contain a relatively high bacterial diversity, which can then contaminate the milk as it is being collected from the teat [4]. Furthermore, even a truly aseptically-collected sample can be contaminated by DNA contained in molecular extraction and library preparation kits, i.e. the so-called “kit-ome” [27–29].

Given the numerous pathways by which exogenous bacteria and/or DNA can contaminate a sample, it is expected that most microbiome datasets will contain some level of non-native DNA sequences. Because of this, many microbiome researchers have developed methods and strategies for differentiating exogenous or “contaminating” microbial signatures from the true microbial composition of a sample. Such advancements include additions to study design, inclusion of extensive positive and negative control samples [30]; and use of synthetic DNA to allow for absolute quantification of microbes [31]. However, it is exceedingly difficult to differentiate contaminating versus endogenous sequences because many bacterial taxa are ubiquitous and could indeed be from both contaminants and endogenous sources within the same sample dataset. To address this issue, researchers have leveraged statistical distributions to identify sequences that likely originated from contaminating cells or DNA [32]. However, many of these advancements have not yet been fully leveraged for milk microbiome research [28,33]. This gap is particularly important because previous studies have reported that a large proportion of 16S rRNA sequences are “shared” between various udder samples and stripped milk samples [24,33], highlighting the necessity of accounting for components within the various niches of the udder to understand their contributions to the stripped milk microbiome. Additionally, understanding the origin of sequences in a microbiome dataset is critical for interpretation, for example to understand potential mechanisms that underlie observed epidemiological associations between microbiome dynamics and animal health. Furthermore, it is nearly impossible to develop therapeutic or preventive cow and herd health strategies if the origin of microbes is ambiguous, i.e., whether they originated from the milk itself, from the external or internal epithelium, from the ambient air in the milking parlor, or from lab-based sample handling procedures. To advance dairy microbiome research, we must address the current ambiguity in the literature regarding the “true” versus “contaminating” milk microbiome.

Therefore, the aim of this study was to obtain an accurate description of the true microbial status of the different compartments of the udder and mammary gland in dairy cows without mastitis. To achieve this aim, we used aseptic sampling techniques and positive as well as negative controls during sample collection and processing. Negative controls were used to identify likely bacterial contaminants and their sources using two available statistical algorithms. We hypothesized that (1) the bovine mammary gland contains a “microbiome gradient” that imparts an increasingly diverse milk microbiome as the milk moves from mammary alveoli through ducts into the gland cistern and internal teat canal, and that (2) current milk sampling techniques artificially inflate the microbial diversity of stripped milk by introducing microbes from the external teat epithelium and ambient air.

## Materials and methods

### Study design and animals

This cross-sectional study utilized dairy cows from the University of Minnesota Dairy Cattle Teaching and Research Center. In total, 90 Holstein dairy cows from the herd were assessed for study enrollment on January 6^th^, 2021. Only cows that met the following eligibility criteria were enrolled; in their first, second or third lactation, >90 days in milk at the time of enrollment; a somatic cell count (SCC) of < 200×1,000 cells/mL, and no history of clinical mastitis in the last 30 days. Twenty-three cows met these requirements from which 14 cows were randomly selected for enrollment into the study. These 14 cows on average were 182 days in milk (DIM, range from 92 to 456 DIM), had a milk yield of 38 kg/d (range from 15 to 50 kg/d), and a milk SCC of 91×1,000 cells/mL (range from17 to 325×1,000 cells/mL) on the sampling day (Table S1). From each cow, teat apex, teat canal, stripped milk and gland cistern milk samples were collected for microbiome analysis (Figure 1A). In addition, air and sampling blanks were collected in the milking parlor to identify potential microbial contaminants in the downstream sequence data analysis (Figure 1C). All the procedures and activities of this study were approved by the University of Minnesota Institutional Animal Care and Use Committee (protocol number 1904-36973A).

**Figure 1.**
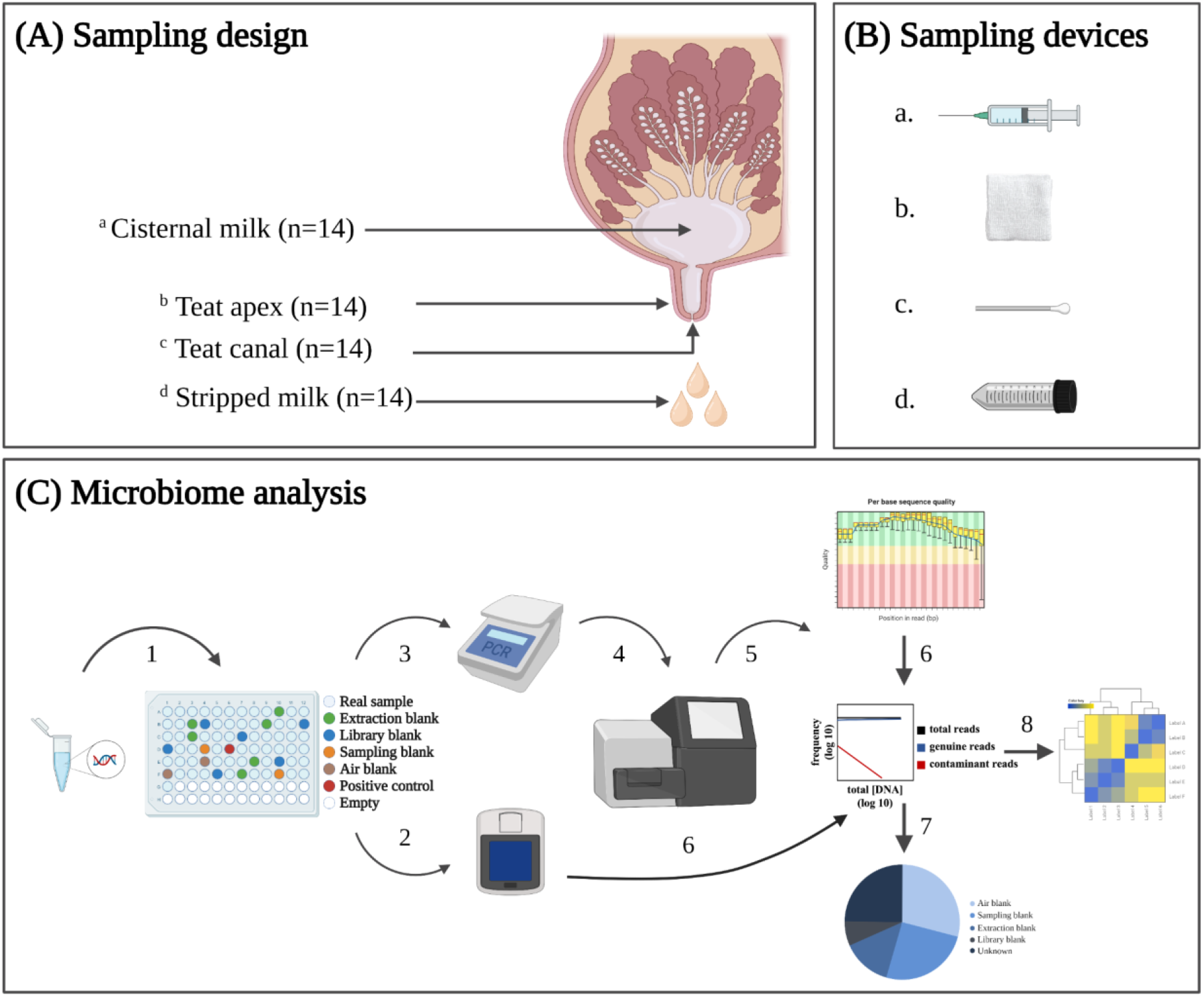
Sampling design (A), sample collection devices (B), and workflow for microbiome analysis (C). Within A and B, lower case letters link sample type with sampling device. Within C; 1) DNA extraction; 2) DNA concentration measurement; 3) qPCR for quantification of the 16S gene; 4) Short-read sequencing of the 16S V4 hypervariable region; 5) Quality control of sequence data; 6) Identification of contaminant sequences; 7) Identification of sequence source samples; and 8) Microbiome analysis. Created with BioRender.com.

### Sample collection procedure

All sampling occurred on the same day. A single hindquarter from each cow was randomly selected for sampling to avoid the potential bias between front and hind quarters. Disposable gloves were used and discarded between each sample type and between each sampled cow. Teat apex and teat canal samples were collected before the milk samples to avoid potential contamination from residual milk that would coat the epithelium of the streak canal after collection of the milk. Teat apex samples were collected first from the distal one-third of the apex, using a sterile, pre-moistened gauze square soaked in germ-free phosphate buffer saline solution (PBS). The gauze square was removed from a whirl-pak bag and gently scrubbed along the teat end and then placed back inside the bag. Teat canal samples were then collected by inserting a sterile cytobrush swab inside the streak canal and rotating the swab 360° before removal and placement inside a whirl-pak bag.

Stripped milk was collected prior to cisternal milk to avoid potential contamination (including blood) flowing from cisternal milk to stripped milk. Before the collection of milk samples, teat skin was prepped with iodine solution and 70% alcohol. After discarding the first three to four streams of milk, stripped milk samples (20 mL) were then collected in sterile falcon tube [23]. The gland was then cleaned with an aseptic wash of iodine solution and 70% alcohol and lidocaine was administered subcutaneously as local anesthetic. Cisternal milk samples were collected by puncturing the mammary gland cistern with a sterile 20-gauge collection needle and the collected milk was released into two 10 mL vacuum collection tubes. Four sampling blanks were also collected as negative controls; “sampling blanks” (N=2) consisted of gauze squares sealed in a whirl-pak bag and not opened until DNA extraction and “air blanks” (N=2) that consisted of gauze squares opened in the sampling location and waved in the air for 30 seconds but not used for actual sampling of the cows. All samples were placed inside a cooler during sampling, and then brought immediately to the University of Minnesota Food Centric Corridor for further processing. Aliquots of 1 mL of stripped milk and 1 mL of cisternal milk were placed inside separate sterile vials in a biosafety cabinet and stored at -20 ℃ until submission for microbiological culture. The remaining milk was stored at -80 ℃, along with the teat apex gauze, teat canal cytobrush, and sampling and air blank samples.

### Bacterial Culture

Aliquots of stripped and cisternal milk stored at -20 ℃ were submitted to the Udder Health Lab at the University of Minnesota for bacterial culture and taxonomic identification within two weeks of their original collection date according to a previous protocol [34]. Briefly, milk samples were plated onto 5% sheep blood agar using a 0.01 mL calibrated loop and incubated in aerobic conditions at 37 ± 2 °C for 42 to 48 h. Samples were classified as contaminated if more than 2 isolates from different bacterial taxa were recovered. Cultured isolates were used as input to a Matrix Assisted Laser Desorption Ionization-Time of Flight (MALDI-TOF) mass spectrometer (Microflex, Bruker Daltonics Inc., Billerica, MA) for taxonomic identification [35]. Peaks produced by each isolate were analyzed by the MALDI-TOF Biotyper reference library using the following confidence level; >2.0, species level classification recorded; 1.8–2, genus level classification recorded; <1.8, MALDI-TOF diagnosis not recorded and traditional identification methods, such as differential growth on selective media, colony morphology and Gram stain, were used for taxonomic classification [35].

### Propidium monoazide treatment

All teat skin and milk samples were subjected to propidium monoazide (PMA) dye treatment to separate DNA from variable and non-viable bacteria [16]. In a 1.5 mL centrifuge tube, 10 µL of PMA solution (1 mg/mL) was added to 1mL of milk sample for a final concentration of 10 µg/mL. Negative controls (N=5) were included during the PMA treatment process, consisting of molecular grade water added to five randomly selected tubes. All solutions were then incubated at room temperature in the dark with agitation for 5 minutes. Samples were then placed on ice and exposed to a 500W halogen light (SMART electrician, Menards, St. Paul, MN) at a distance of 20 cm for 5 minutes with occasional inversions to ensure total exposure. The samples were then centrifuged at 12000 rcf for 5 minutes, after which the supernatants were removed, and the pellet were washed three times with an equal volume of phosphate buffer solution (pH 7.4) to remove any excess PMA before being stored in -20℃ until DNA extraction.

### DNA Extraction, Library Preparation, and Sequencing

All teat skin, teat canal, milk samples, their PMA treatment counterparts, and all sampling blanks were subjected to DNA extraction using the PowerSoil Pro Kit (Qiagen, Cat No. 47016, Hilden, Germany), automated on the QiaCube Connect instrument (Qiagen, Cat No. 9002864, Hilden, Germany). Briefly, whirl-pak bags containing teat apex gauze or teat canal cytobrush samples were thawed for 20 minutes in a bio-safety cabinet sterilized using 70% EtOH. Once thawed, the gauze or the tip of the cytobrush was cut using a metal pair of scissors. The dirtiest part of the gauze or the cytobrush tip was placed inside a PowerBead Pro tube. This process was repeated for each teat apex and canal sample using metal forceps and scissors that were sterilized with a glass-bead sterilizer (250F). Milk samples DNA extraction proceeded according to the manufacturer’s instructions. The instrument was run 6 times, once for each batch of 12 samples that were processed. Each batch included an extraction blank control (“Extraction”, N=6), containing 800 uL of CD1 lysis buffer in an empty bead beating tube without any sample material. The purpose of the extraction blank control was to identify and account for reagent contamination from the DNA extraction kit. A single positive control sample (“Positive”), containing a known concentration and taxonomy of 8 bacterial and 2 fungal species was also included as an internal control (Zymo Research Corp, Cat No. D6310). Extracted nucleic acids from each sample were randomized into a 96-well rack (Figure 1C) and approximately 20 uL of each sample was transferred from the rack to a PCR plate, covered with an adhesive seal, and then submitted to the University of Minnesota Genomics Core (UMGC) for library preparation and sequencing.

Extracted DNA was quantified and assessed for quality by both Nanodrop UV/VIS spectrophotometry (ThermoFisher, USA) and fluorometry using PicoGreen staining (BioTek, USA). The 16S rRNA gene copy number in each sample was determined using qPCR. Libraries were prepared by amplifying the V4 region of the 16S ribosomal RNA gene, using primer Meta_V4_515F: GTGCCAGCMGCCGCGGTAA and Meta_V4_806R: GGACTACHVGGGTWTCTAAT (Gohl et al., 2016). Negative controls for library preparation (“Library”, N=5) were included during the library preparation process, consisting of molecular grade water added to five randomly selected wells. Prepared libraries were sequenced to an expected sequencing depth of 100,000 paired-end reads per sample on an Illumina MiSeq instrument (Illumina Inc, San Diego, CA) using a 600 (2×300 base pair) cycle reagent kit (Illumina Inc, San Diego, CA).

### Bioinformatics and sequencing efficiency

All the following bioinformatic and statistical analyses were performed in R Statistical Software (v4.1.2; R Core Team 2021; https://www.r-project.org/) and plots were generated using the ggplot2 v3.4.2 package for R [36]. Raw sequence data were processed through the DADA2 (v1.22.0) pipeline for quality filtering, denoising, and microbial community inference [37]. The *filterAndTrim* function was used to quality filter the raw sequence data. Primer pairs were removed by trimming the first 20 and 17 base pairs from the 5’ ends of forward and reverse reads, respectively. Forward and reverse reads were truncated to a length of 270 and 200 base pairs based on the observed distribution of quality score and sequence reads containing ambiguous base pairs and PhiX were discarded. Forward and reverse reads with a maximum expected error rate greater than 3 or 4 base pairs, respectively, were also discarded. The *learnErrors* function was used to estimate the expected error rates produced by the Illumina MiSeq sequencer. Error corrected reads extending beyond or below the expected length of the sequenced amplicon were discarded. Forward and reverse reads were concatenated using the *mergePairs* function. Merged reads were used as input to the *removeBimeraDenovo* function to identify and remove chimeric sequences. The merged reads with a length beyond 250 and 258 base pairs were discarded based on the distribution of sequence reads and the length of expected V4 region. Chimera-free amplicon sequence variants (ASVs) were aligned to the SILVA reference database (v138.1) for taxonomic assignment using the *assignTaxonomy* function. The ASV abundance matrix, taxonomy table and sample metadata were used to generate a phyloseq object for microbiome data analysis using the *phyloseq* (v1.38.0) R package [38].

The sequence data for the positive control sample was aligned to the manufacturer’s sequence database (https://s3.amazonaws.com/zymo-files/BioPool/ZymoBIOMICS.STD.refseq.v2.zip) containing 16S rRNA reference sequences for each mock bacterium, using the same procedures described above. Extraction and sequencing efficiency were measured by inspecting the distribution and taxonomy of the positive control sample. Sequence features classified to *Listeria*, *Pseudomonas*, *Bacillus, Escherichia*, *Salmonella*, *Lactobacillus*, *Enterococcus*, and *Staphylococcus* spp. were extracted and their distribution visually assessed via bar plots.

Due to the lack of classification for most reads in the dataset of PMA-treated samples, all the following microbiome analyses were performed on the dataset generated by samples without PMA treatment. We further applied BLAST (https://blast.ncbi.nlm.nih.gov/Blast.cgi) to align the most abundant reads in the dataset of PMA-treated samples to all nucleotide collections using default settings [39].

### Identification and removal of potential contaminants

Potential contaminants obtained during sampling, DNA extraction and sequencing were identified using decontam [40] and SourceTracker [41] packages in series. First, the phyloseq object was subjected to the decontam package (v1.14.0) to identify potential sequence contaminants introduced during DNA extraction and library preparation. The *isContaminant* function was used to identify contaminants using both the frequency- and prevalence-based methods at the threshold of 0.5 after plotting the distribution of score statistics, as recommended by Davis et al [40]. Sequence features classified as potential contaminants using the frequency method were visualized by plotting the abundance of the contaminant within each sample as a function of that sample’s measured 16S qPCR concentration using the *plot_frequecy* function. The prevalence method was run two times, using extraction blanks and library preparation blanks as negative controls, respectively, since different contaminants were expected from each type of blank. ASVs identified as contaminants were then removed from the phyloseq object.

The contaminant-subtracted phyloseq object was then subjected to the SourceTracker package to identify the proportion of ASV counts in the animal samples (i.e., teat apex, teat canal, stripped milk and cisternal milk) that could be attributed to negative controls. In this analysis, negative controls were defined as source environments and animal samples were defined as sink environments. The proportion of each individual ASV arising from each source was then predicted and removed from the animal samples according to previously reported methods [32]. The reduced dataset was then used for all subsequent microbiome analyses. The SourceTracker analysis was then used to understand the contribution of 1) the negative controls, teat apex and canal (sources) to the cisternal milk (sink) and 2) the negative controls, teat apex, canal and cisternal milk (sources) to the stripped milk (sink).

### Microbiome Diversity Analysis

To evaluate the prevalence of potential mastitis pathogens in the animal samples, we subsetted the genus-level count matrix of the 16S rRNA data to only include the potential pathogen candidates reported in a previous study [42]. Alpha diversity of true samples and negative controls was estimated by calculating richness, Shannon diversity index and inverse Simpson’s index of diversity, using the *estimate_richness* function in the phyloseq package. Alpha diversity metrics stratified by sample types were visualized via raincloud plot using the gghalves (v0.1.4) package [43].

Beta diversity was estimated by computing pairwise Bray-Curtis distances between all animal samples and negative controls, and microbiome dissimilarities were visually compared by non-metric multidimensional scaling (NMDS) using the *ordinate* function in the phyloseq package. Ordination fit was assessed by increasing the “trymax” parameter i.e., the maximum number of random starts used to search for a stable solution, until stress value <0.2 was obtained. The proportion of variation in the microbiome that was due to sample type was estimated using a permutational multivariate analysis of variance (PERMANOVA) [44]. The *pairwise*.*adonis2* function implemented in the vegan package was used after 999 permutations [45]. The homogeneity of variance between sample types was estimated by the *betadisper* function implemented in the vegan package and ANOVA.

### Statistical analysis

To describe the difference in sequencing depth, 16S gene copies (log10) and alpha diversity indices between different sample types, a generalized linear model was fit using the *glm* function. Model significance was assessed using ANOVA. Adjusted means were computed between sample types using the emmeans package [46], with significance being determined at the 0.05 level.

## Results

### Bacterial culture

Cisternal and stripped milk samples were subjected to standard microbiological culture typically used for mastitis testing (Table 1). Among cisternal samples, one sample exhibited growth of *Staphylococcus chromogenes*, while the remaining 13 exhibited no bacterial growth. Among stripped milk samples, 7 exhibited no bacterial growth, 3 exhibited growth of *Corynebacterium sp.*, 2 of gram-negative bacteria, 1 of *Staphylococcus chromogenes*, and 2 were contaminated (i.e., growth of three or more bacteria). The cisternal milk sample with growth of *Staphylococcus chromogenes* was collected from one of the cows (ID, 3019) whose stripped milk sample also grew *Staphylococcus chromogenes* and was classified as contaminated due to the growth of more than two microorganisms.

**Table 1.**
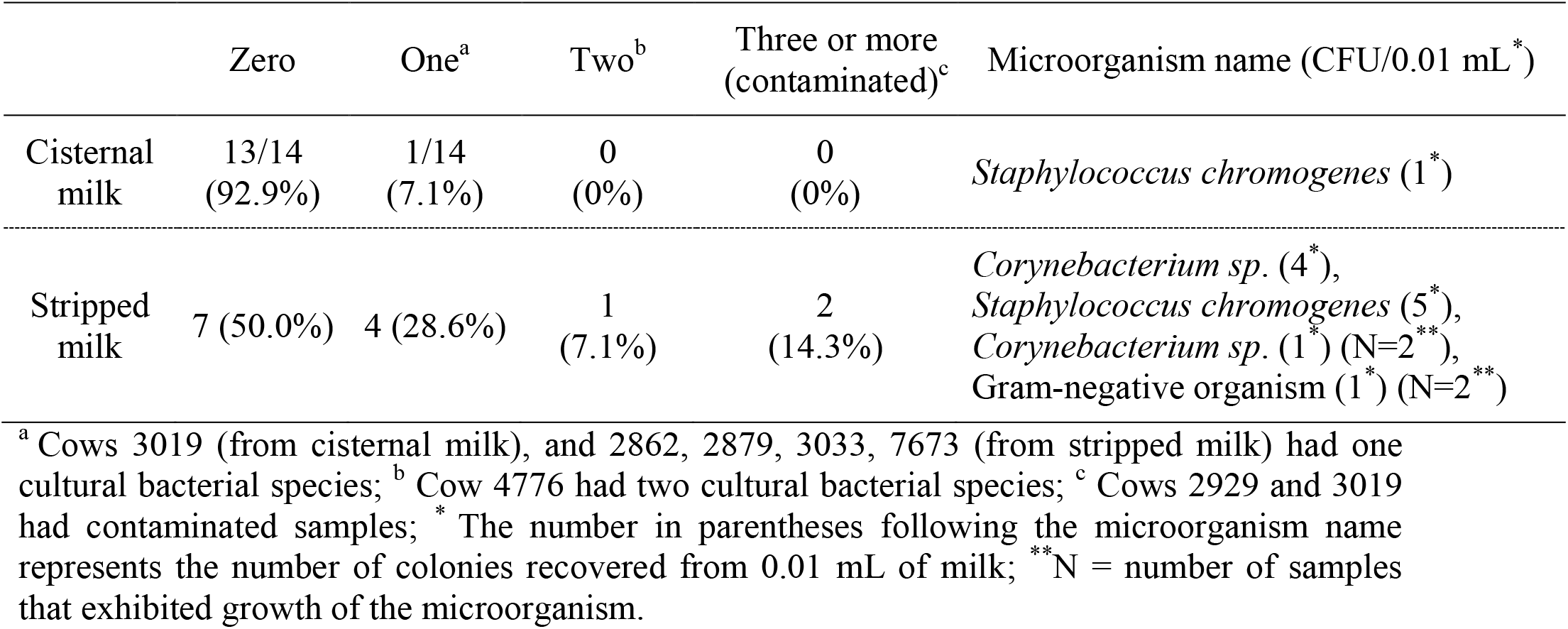
Number (%) of samples containing zero, one or two distinct microorganisms, stratified by the type of milk sample collected (cisternal versus stripped).

### Sequencing depth and quality

In total, 10.8 M raw paired-end reads were generated from 72 samples, including animal samples and negative controls, with an average of 149,654 reads per sample (range: 563- 269,313). A total of 8.5 M remained after quality filtering and merging the forward and reverse sequence reads, and 6.9 M paired-end sequence reads remained after removing chimeras. One stripped milk sample (Cow ID: 4778) was removed from further analysis because it contained a lower number of reads than the extraction blanks, suggesting a failed library. After removing this outlier sample, no significant differences in sequencing depth were observed among sample types, including animal samples (i.e., teat apex, teat canal, stripped milk and cisternal milk), sampling controls (i.e., air and blank) and extraction controls (Figure 2A, ANOVA, *P* > 0.05). Library controls showed a significantly (ANOVA, *P* < 0.05) lower number of reads than the other samples.

**Figure 2.**
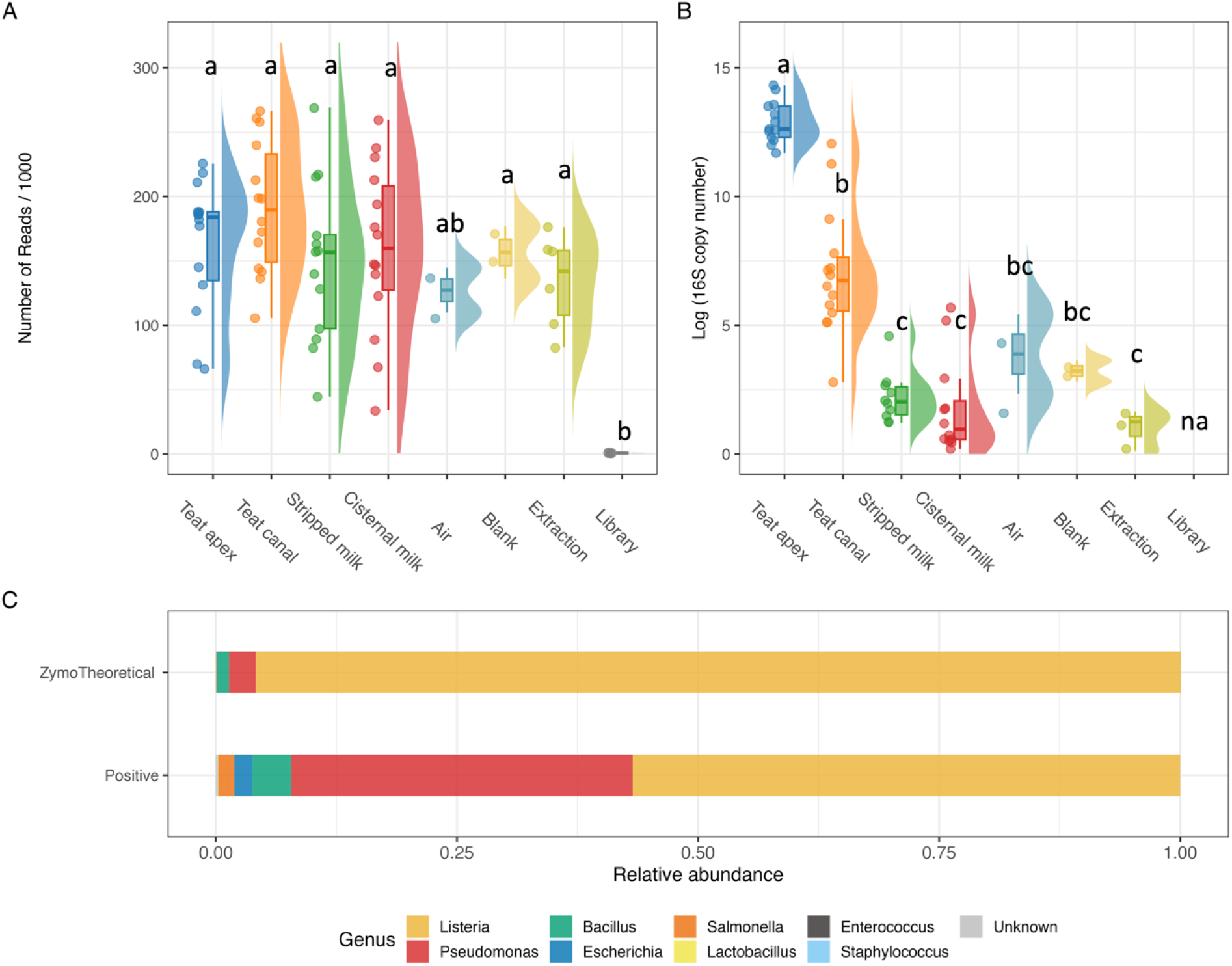
(A) sequencing depth and (B) 16S gene copy number across sample types, including animal samples and negative controls. (C) relative abundance of genus composition in positive control sample and Zymo Theoretical (theoretical standard) samples, taxa that are not matching any of the mock community taxa were grouped into “Unknown”. Different letters within each panel indicate significant differences between sample types. na = not applicable.

The log-transformed 16S copy number was significantly higher in teat apex samples than all other sample types (ANOVA, *P* < 0.05), followed by teat canal samples, which were similar to sampling controls (i.e., air and blank) (Figure 2B). Stripped milk and cisternal milk samples had the lowest 16S copy number among animal samples, and these values were not significantly different from sampling and extraction controls (Figure 2B).

The positive control sample generated 235K raw sequences, and the 186K sequences that remained after quality control were assigned to 34 taxa using the Zymo reference database (Figure 2C). Six of these 34 taxa were from the mock community, while the rest were unidentified but accounted for 0.2% of the total sequences generated from the mock community sample. *Escherichia* and *Staphylococcus* (theoretical abundances of 0.061 and 0.01%, respectively) were not recovered in the positive control sequence data, possibly due to their low abundance within the mock community. In addition, *Pseudomonas* was overrepresented with 35.4% of the sequence data (theoretical relative abundance 2.8%), while *Lactobacillus* was underrepresented at 56.8% (theoretical relative abundance 95.9%).

### Decontamination

Decontamination was first performed using decontam by both prevalence and frequency methods (Figure 3). The score statistic assigned by decontam was used to classify sequence features (ASVs), with a score <0.5 indicating a contaminant. Using the frequency method, the majority of sequencing features (prevalence > 2) were assigned scores close to 1.0, suggesting that they were not contaminants (Figure 3A). Using the prevalence method, we observed a unimodal distribution of scores around 0.5 using either library blanks (Figure 3B) or extraction blanks (Figure 3C) as negative controls. The contaminants identified by the prevalence method, using both library and extraction as negative controls, confirmed that true contaminants (teal, under the diagonal line) had higher prevalence in negative controls than in true samples (Figure 3D).

**Figure 3.**
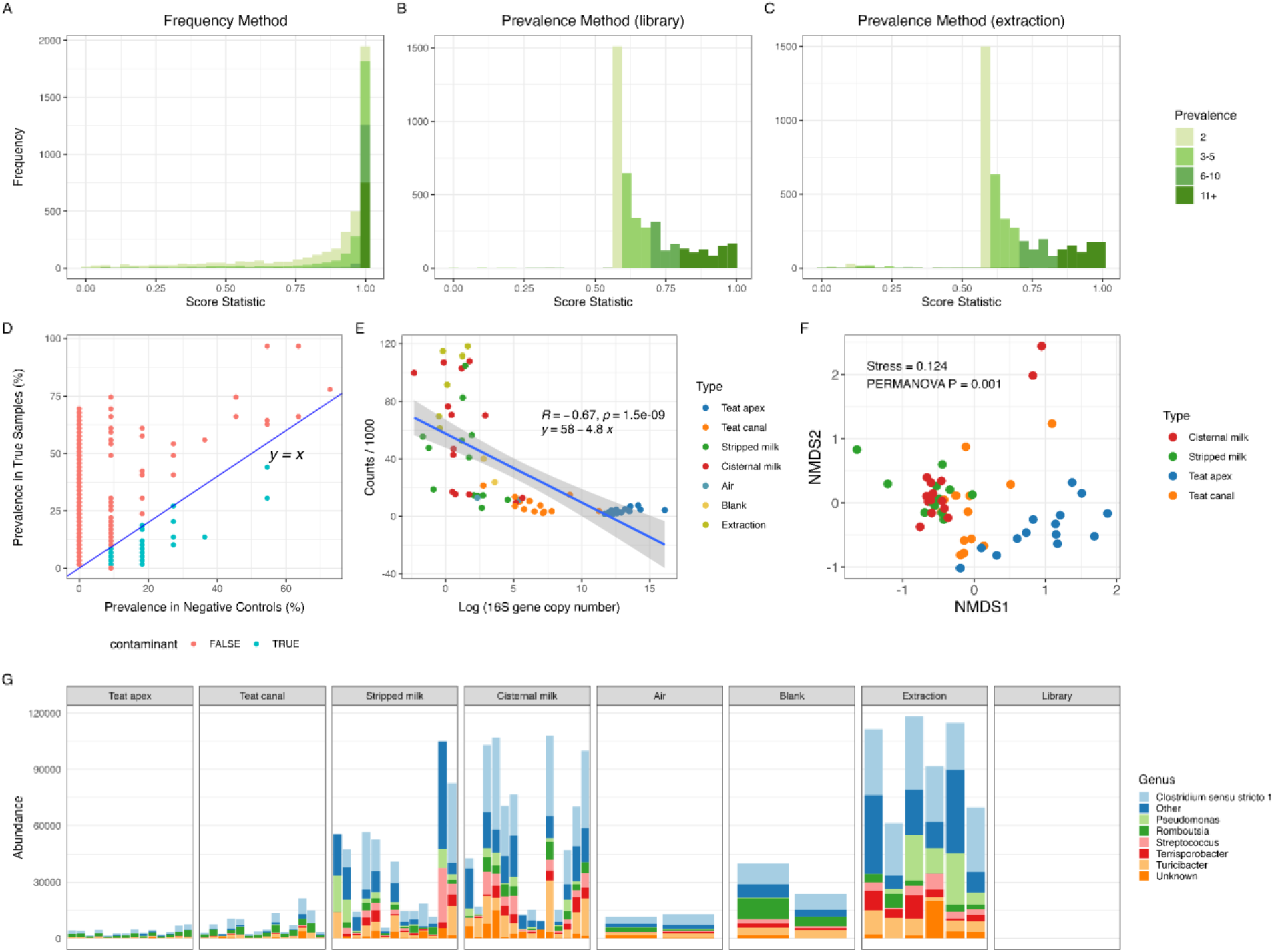
Contaminants identified using decontam [40]. Histograms showing the distribution of score statistics assigned to amplicon sequence variant (ASV) present in two or more samples using (A) frequency method, (B) prevalence method with library blanks as negative control, and (C) prevalence method with extraction blanks as negative control. Color intensity indicates the number of samples in which each ASV was present (i.e., prevalence). (D) prevalence (%) of contaminants (teal, “TRUE”) and non-contaminants (pink, “FALSE”) identified by the prevalence method in true animal samples (y-axis) and negative controls (x-axis), (E) correlation between 16S copy number (x-axis) and counts of contaminants/1000 (y-axis) from each sample identified by the combined frequency and prevalence method, (F) NMDS ordination of Bray-Curtis distances of *decontam* contaminants identified from milk, teat apex and teat canal samples, (G) genus-level counts of contaminant reads (“Abundance”, y-axis) identified by combined frequency and prevalence methods; genera with relative abundance <5% and prevalence <20% across all samples were grouped into “Other”.

The frequency method identified 422 contaminant ASVs, while the prevalence method identified 11 and 111 contaminant ASVs using library blanks and extraction blanks as negative controls, respectively. In total, 516 ASVs representing 2.1M reads were identified as contaminants using both methods, which accounted for 31.9% of the filtered reads. Among the sample types, extraction blanks contained the highest abundance of contaminants, followed by cisternal milk and stripped milk (Figures 3E and 3G). Teat apex and teat canal samples contained a lower abundance of contaminants compared to sampling controls (i.e., air and blank). The number of reads originating from contaminant ASVs was inversely correlated with 16S gene copy number, suggesting that low biomass samples were prone to microbial contamination during sample collection and processing (Figure 3E). The structure of sequence features within the contaminant data showed significant differences between sample types (Figure 3F, PERMANOVA, *P* = 0.001, R^2^ = 0.32). Only the contaminants identified in stripped milk and cisternal milk showed no significant differences in microbial composition based on beta-diversity analysis, suggesting that the profile of contaminants in these samples was similar (Table S2, PERMANOVA, *P* = 0.200, R^2^ = 0.049). Few contaminants were identified in library controls using decontam because they were used as a negative control for the prevalence method, and they did not have corresponding 16S copy number information for inclusion in the frequency method. *Clostridium sensu stricto 1* was dominant among identified contaminants across all sample types, while *Pseudomonas*, *Romboutsia*, *Streptococcus*, and *Turicibacter* also showed relatively high abundance (abundance >5%) and prevalence (>20%) amongst the contaminant data (Figure 3G). All identified contaminant reads were removed from the dataset and remaining reads were subjected to SourceTracker to identify the proportions of contribution from defined sources.

After running decontam, the proportion of reads in each sample originating from negative controls (i.e., air, blank, extraction, library) was calculated using SourceTracker (Figure 4A). A large proportion of reads within the samples from the teat apex (66.2%) and teat canal (57.8%) were sourced from negative controls (primarily sampling and air blanks), while these sources represented only a small proportion of the reads in the cisternal milk and stripped milk samples (13.0% and 23.3%, respectively; Figure 4A). Library blanks showed little contribution to any of the milk, teat apex or teat canal samples suggesting that the library preparation process introduced little contamination to true samples (Figure 4A). Across all the cisternal milk samples, the main sources of contaminating reads were from extraction blanks (7.6%), but rarely from air blanks (1.5%) and sampling blanks (1.8%). This differed from stripped milk samples where air blanks comprised the highest source of contamination, followed by sampling blanks and extraction blanks (11.7%, 5.1% and 4.7%, respectively). It is notable that the contamination sources for the milk samples were highly variable across individual samples (Fig 4A-B), such that each sample seemed to have its own dominant contamination source. This highlights the unpredictability in defining sources that contribute to the microbiome data generated from bovine milk samples. On the other hand, teat apex and teat canal showed a consistent pattern of contamination sources, which were mainly air blanks (mean = 38.3%, SD = 19.1%) and sampling blanks (mean = 23.2%, SD = 13.2%). Most of these identified contaminants were from genera *Corynebacterium*, *Acinetobacter*, *Bacteroides*, *UCG-005*, *Kocuria*, and *Methanobrevibacter* (Figure 4C). Sample type had a significant effect on the microbial composition of sourced contaminants identified by SourceTracker (Figure 4D, PERMANOVA *P* = 0.001, R^2^ = 0.24). However, there were no significant differences in the microbial community of contaminants between stripped and cisternal milk samples (PERMANOVA *P* = 0.222, R^2^ = 0.048) or between teat apex and teat canal samples (PERMANOVA *P* = 0.353, R^2^ = 0.039) (Table S3). When teat apex and teat canal samples were included in potential sources of contaminants in the milk samples, SourceTracker indicated that a majority of the stripped milk samples contained a high proportion of sequences from the teat apex and teat canal, while only one of the cisternal milk samples (Cow ID, 3019) was dominated by contaminants originating from the teat canal (Figure 4B).

**Figure 4.**
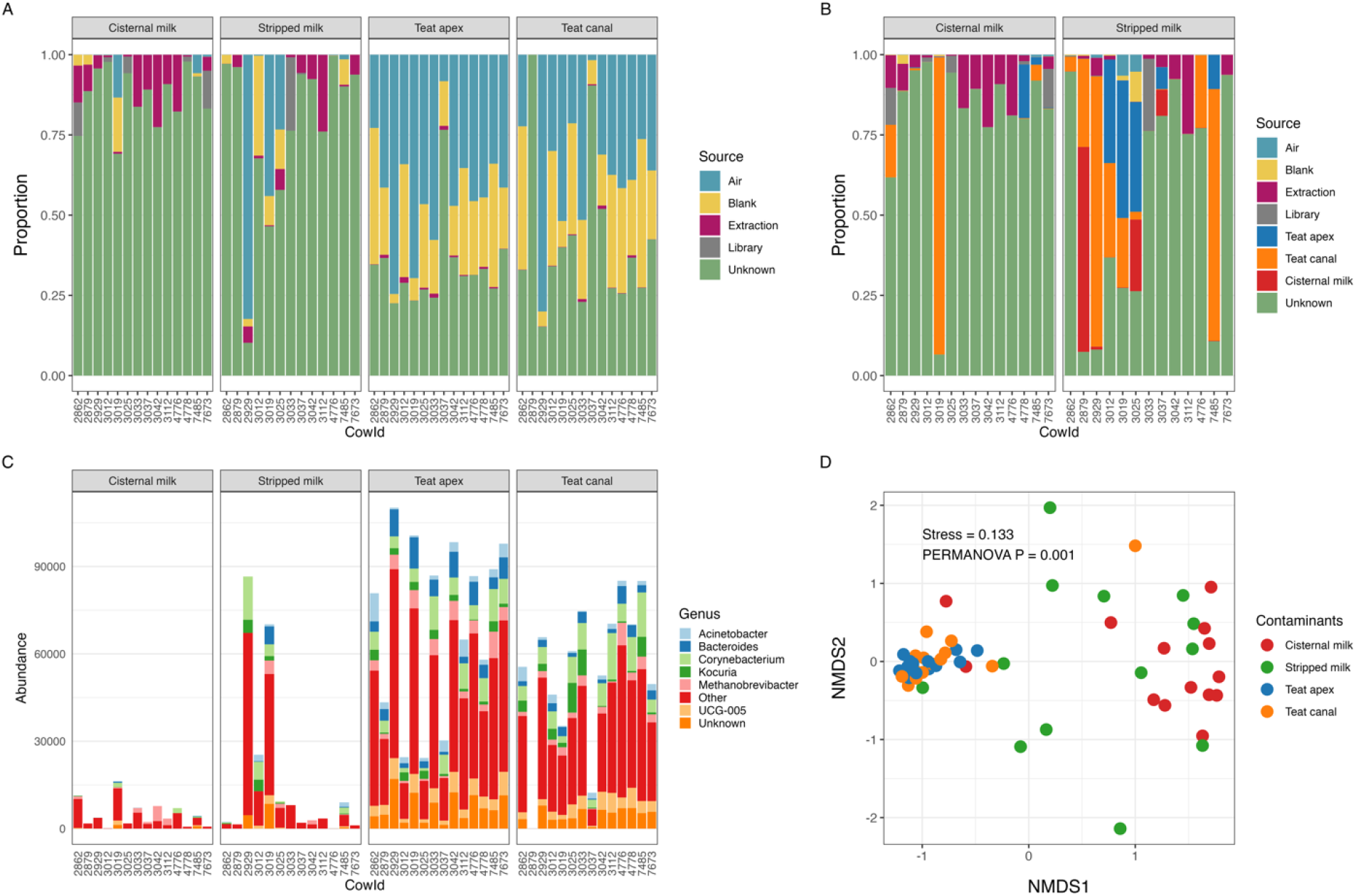
Source of contaminants identified by SourceTracker. (A) proportion of reads identified as contaminants using negative controls (i.e., air, blank, extraction, library) as sources; (B) proportion of reads identified as contaminants using negative controls, teat apex and teat canal as sources; (C) counts of genus-level contaminant reads using negative controls as source environments; genera with relative abundance < 5% and prevalence < 25% across all samples were grouped into “Other”; (D) NMDS ordination of Bray-Curtis distances of SourceTracker contaminants identified in milk, teat apex and teat canal samples.

### Microbial diversity

We next assessed how quality filtering, decontam and SourceTracker impacted alpha diversity metrics. Across all samples, a relatively high proportion of sequencing reads were removed during the QC process (Figure 5A). After removal of putative contaminants by both decontam and SourceTracker, the samples within each sample type retained only ∼25% of the total reads generated. Identification of contaminants differed by sample type, with decontam identifying most of the contaminants in cisternal and stripped milk samples (36% and 27%, respectively) and SourceTracker identifying most of the contaminants in the teat apex and teat canal samples (44% and 29%, respectively; Figures 5A and S1). Reads identified as contaminants by decontam represented a fairly consistent number of unique ASVs (N = 26-32) across sample types. In contrast, reads identified as contaminants by SourceTracker belonged to a very large number of unique ASVs within the teat apex and teat canal samples (534 and 396, respectively) and a much smaller number of unique ASVs in the cisternal and stripped milk samples (10 and 21, respectively). Patterns of alpha diversity (as measured by microbial richness, Shannon diversity and Inverse Simpson) remained relatively stable across the decontamination process (Figure 5B-D). At each step of the decontamination process, the order of diversity (from highest to lowest) was teat apex > teat canal > air > blank > stripped milk > cisternal milk > extraction > library. In all cases, the teat canal and teat apex samples showed significantly higher richness and Shannon’s index than cisternal and stripped milk samples. However, across all alpha diversity indexes, teat apex samples did not have significantly different diversity than teat canal samples, and the diversity of both teat apex and teat canal samples was similar to that of the sampling controls (i.e., air and sampling blanks). Similarly, stripped milk and cisternal milk had no significant differences in all alpha diversity indices across all steps of the decontamination process, and their diversity values were similar to that of extraction and library blank controls. Thus, relative ranking of sample types by their alpha diversity index was teat apex = teat canal = air = blank > stripped milk = cisternal milk = extraction = library.

**Figure 5.**
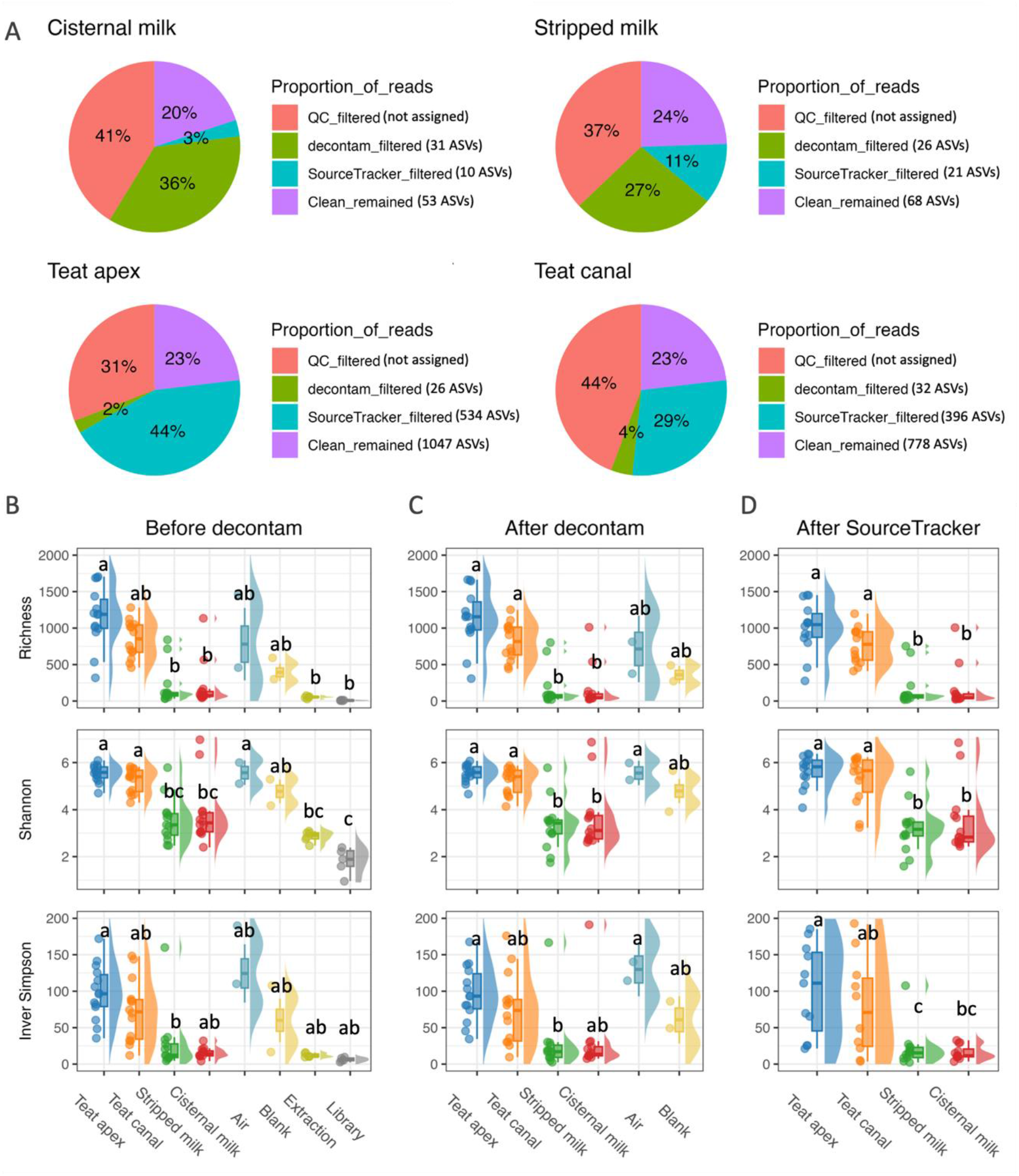
Alpha diversity before and after decontamination with decontam and SourceTracker. (A) Pie charts showing the percentage of reads that were removed during quality control (“QC”, low quality reads), “decontam”, and “SourceTracker”; as well as the percentage of remaining reads at the end of the decontamination process (“Clean”); stratified by sample type. For each step of the decontamination process, the number in the parentheses indicates the median number of ASVs removed by decontam and Sourcetracker; and the median number of ASVs that remained in the “Clean” reads. Raincloud plots for alpha diversity indices (i.e., richness, Shannon’s, and Inverse Simpson’s) (B) before decontam, (C) after removing contaminants identified by decontam, and (D) after removing contaminants identified by SourceTracker; stratified by sample type. Sample groups with different superscripts had significantly different alpha diversity values within each facet B-D.

Non-metric multidimensional scaling (NMDS) ordination and PERMANOVA were performed to visualize and compare the composition of the microbial communities between different datasets (i.e., before decontam, after decontam, and after SourceTracker) and between sample types (i.e., teat apex, teat canal, cisternal milk and stripped milk) (Figure S2). Results showed that microbial composition at the ASV level varies within the overall dataset and within sample type (PERMANOVA *P* < 0.001 in both cases). Hence, the comparison of microbial community composition was performed on datasets stratified by decontamination step. Subsequent NMDS analysis showed that the microbial composition clustered according to sample type, and this clustering pattern was maintained across the decontamination process (Figure 6A-C). However, milk samples displayed significantly higher beta-dispersion than teat apex and teat canal samples (ANOVA *P* < 0.01), and thus we also stratified by sample type (Figures 6D-I). At each step of the decontamination process, no significant differences in beta diversity were observed between teat apex and teat canal samples or between milk samples (stripped milk vs cisternal milk) (Figures 6 D-I and Table S4). However, teat apex and canal samples and milk samples displayed significant differences in the microbial composition across the decontamination process.

**Figure 6.**
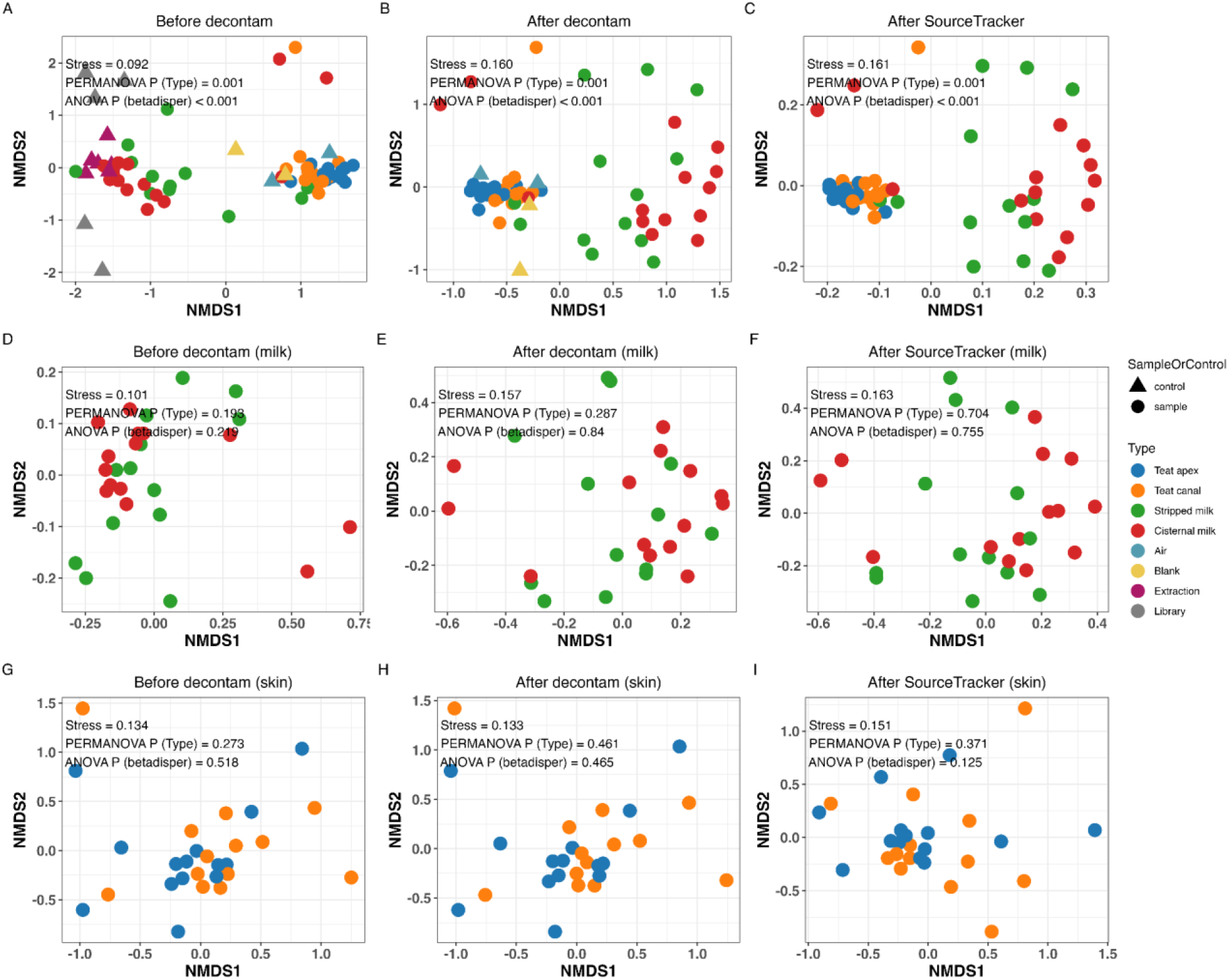
NMDS ordination based on Bray-Curtis distance matrix, using the entire datasets (A) before decontam, (B) after decontam and (C) after SourceTracker, (D-F) using the milk samples datasets, (G-I) using teat apex skin and teat canal sample datasets. PERMANOVA *P* is the *P* value for sample types; ANOVA *P* is the *P* value for betadisper variance between sample types.

### Microbial composition

The microbial composition and relative abundance of top taxa from skin and milk samples shifted across the decontamination process, at both the phylum (Figure 7A) and genus (Figure 7B) levels. After use of SourceTracker, Firmicutes was the dominant phyla across all skin and milk samples, accounting for 41.9% of sequence counts (Figure 7A). Bacteroidota (19.9%), Proteobacteria (13.9%), and Actinobacteriota (13.0%) were also predominant. At the genus level, no clearly predominant microorganisms were observed in either skin or milk samples (Figure 7B), which exhibited remarkable consistency in microbial profiles, except for a few samples with a high relative abundance of *Staphylococcus*. After decontamination, *Staphylococcus* (4.9%) and *Acinetobacter* (4.2%) were the top two genera in relative abundance across all samples. Among the top genera, *Acinetobacter, Bacteroides*, and *Corynebacterium* exhibited higher abundance in skin versus milk samples, while *Clostridium sensu stricto 1* showed the opposite trend. The relative abundance of *Clostridium sensu stricto 1* decreased substantially during the decontamination process, especially in milk samples, demonstrating its importance as a major source of contaminating microbial DNA. *Staphylococcus* showed high relative abundance in cisternal milk (7.9%), stripped milk (5.9%), and teat canal (5.5%) samples, but lower relative abundance in teat apex (0.6%) samples.

**Figure 7.**
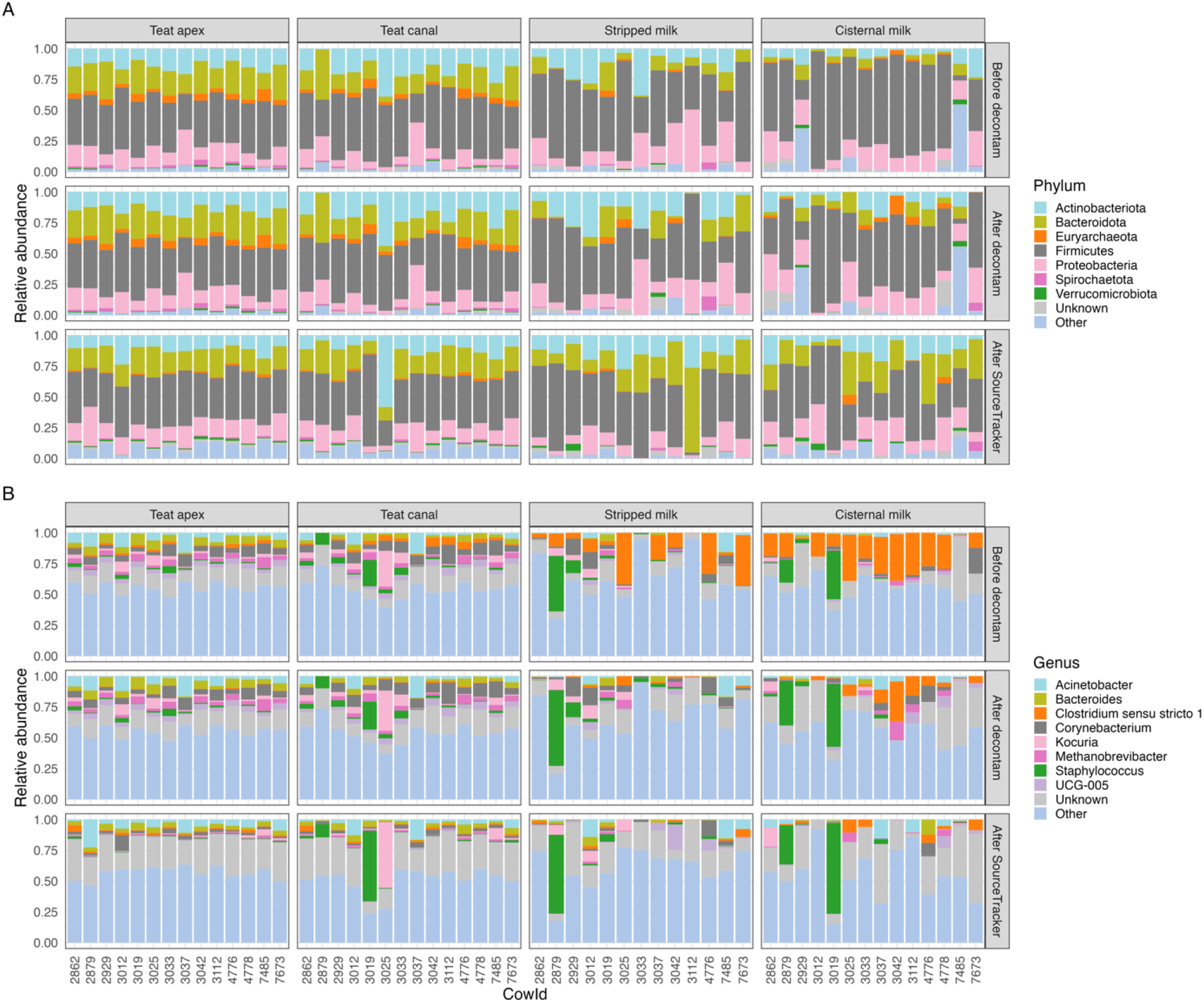
The relative abundance of most abundant taxa in animal samples detected before decontam (top row), after decontam (middle row), and after SourceTracker (bottom row) at (A) phylum level (abundance < 1% and prevalence < 20% were grouped as “Other”) and (B) genus level (abundance < 3% and prevalence < 20% were grouped as “Other”).

### Presence of potential mastitis pathogens

The presence of potential mastitis pathogens in relative abundance across all samples is shown in Figure 8. On average, sequences from potential mastitis pathogens accounted for 14.8% of the reads across all sample types. At the genus level, the potential pathogens with the highest relative abundance were *Staphylococcus* (5.1%), *Acinetobacter* (4.0%), *Corynebacterium* (2.2%), *Aerococcus* (1.7%), *Streptococcus* (1.1%) and *Pseudomonas* (0.7%). Samples from two cows (Cow IDs, 2879 and 3019) had distinctly greater abundance of *Staphylococcus* in their cisternal milk, stripped milk and teat canal samples (Figure 8) and culture results (Table 1), which confirmed the presence of *Staphylococcus* in these milk samples. Interestingly, the milk samples from these two cows also yielded growth of *Staphylococcus* during bacteriological testing. *Streptococcus* sequences also showed high relative abundance in cisternal milk (1.7%), stripped milk (1.2%) and teat canal (1.1%) samples, but only 0.4% relative abundance in teat apex samples. Teat apex contained a high relative abundance of *Acinetobacter* (7.0%) when compared to other sample types.

**Figure 8.**
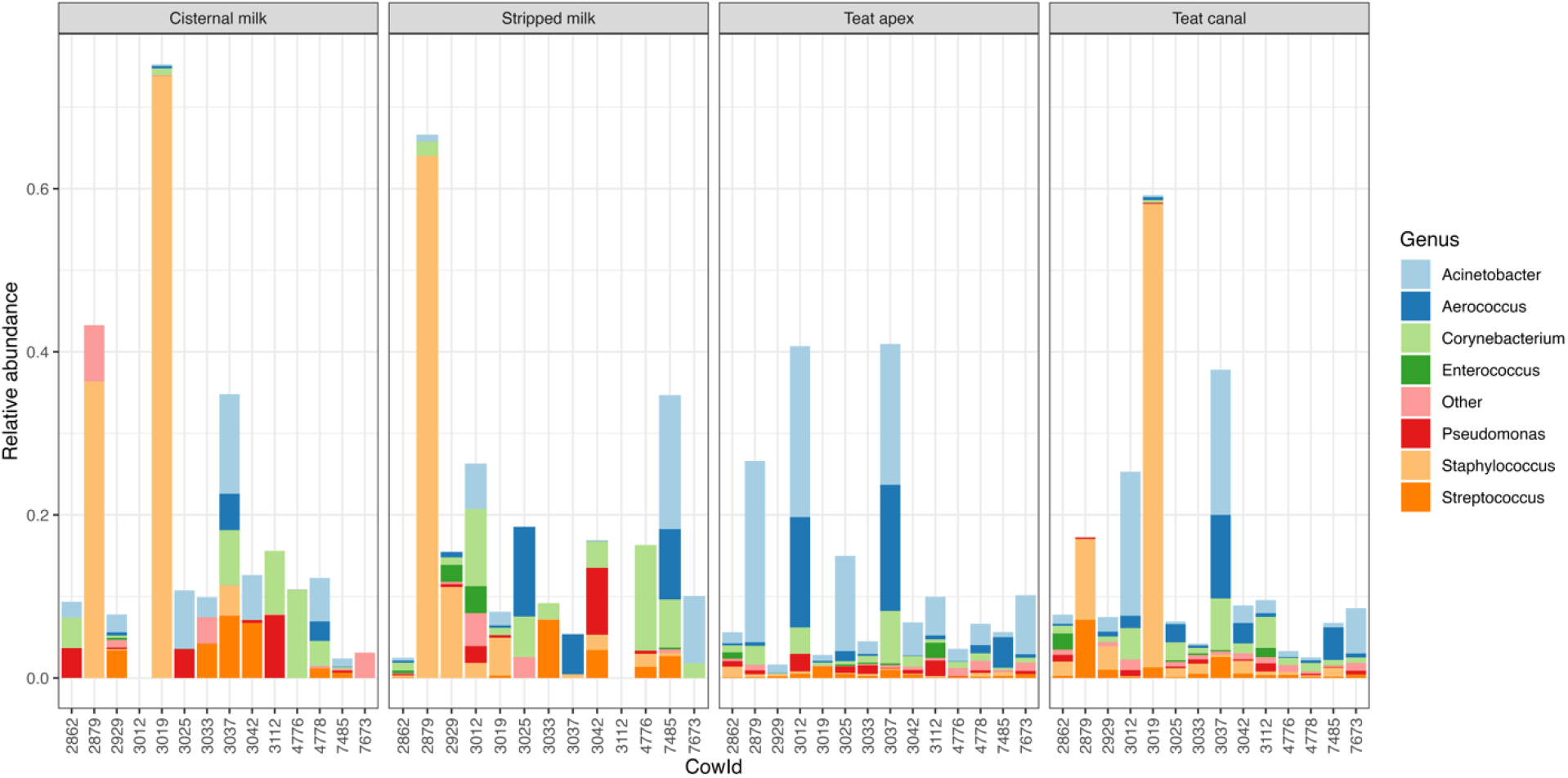
Barplot of the relative abundance (y-axis) of potential mastitis pathogens (proportion of all genus-level counts) in each sample (x-axis), grouped by sample type. Only genera with a relative abundance > 0.1% relative abundance and prevalence > 25% are depicted as individual colors within the bars; the rest are grouped together as “Other”.

### Microbial composition of PMA-treated samples

Sequencing of samples treated with PMA (N=60) generated 1.9M raw paired-end reads and only 0.3M reads remained after removing chimeras. Among these non-chimeric reads, 18.9% were classified as Proteobacteria and 1.58% were assigned to Cyanobacteria at the phylum level (Figure S3A). At the family level, only 7.1% of the non-chimeric reads were classified, and all of them were classified as *Mitochondria* (Figure S3B). Zero non-chimeric reads were classified at the genus level (data not shown). Due to the lack of classification for most reads in the dataset of PMA-treated sample dataset, we performed additional analysis on the top 10 most abundant of these unclassified ASVs. Specifically, we used BLAST to align these reads to the NCBI GenBank [39]. These results all yielded high matches to the *Bos taurus* genome, suggesting that these ASVs originated from intact host cells in the samples. We note here that even the samples from the two cows with culturable *Staphylococcus* did not yield classifiable sequencing reads in the PMA-treated samples dataset.

## Discussion

Contamination is ubiquitous in microbiome studies and is especially problematic for samples with low biomass such as bovine milk. In this study, we aimed to characterize the “true” dairy cow milk microbiome from different compartments of the mammary gland by identifying potential contamination of milk from the external and internal teat epithelial microbiomes and from the sample handling process including sample collection, DNA extraction and library preparation. Using multiple sampling controls and a two-step sequential statistical algorithm approach, we found that the majority of sequence reads were identified as contaminants and thus removed from microbiome analysis. After this contaminant identification and removal process, only 20-24% of the reads remained for microbiome analysis, suggesting that the vast majority of sequence data typically generated from bovine milk and udder samples may represent contaminating microbes and/or DNA. This finding is important both for interpreting previous milk microbiome studies and for designing future studies. Previous results that do not incorporate extensive use of negative controls should be interpreted cautiously, because the majority of the data used in those studies could represent contamination. For future studies, it is important to include numerous negative controls so that the impact of contamination can be quantified and fully appreciated in interpreting results.

The contaminants that we identified in the milk samples mainly originated from the DNA extraction procedure (sometimes called the “kit-ome”). Contaminants of skin samples were mostly consistent with those identified in the sampling devices which indicates these contaminants were primarily from microbes in the air of the milking parlor and laboratory and from gloves worn during handling the devices. After removing all potential contaminant sequences, the milk microbiome profiles were less diverse, more dispersed, and compositionally distinct as compared to the skin microbiome profiles. However, our results indicate that a portion of the stripped milk microbiome might originate from microbes that inhabit the teat apex and teat canal. Even after the statistical decontamination procedures, potential mastitis pathogens were detected in both milk and in skin samples and with relatively high abundance was detected in milk from cows that exhibited no signs of clinical mastitis, which has the potential of developing mastitis.

### Potential sources of contamination in bovine milk microbiome data

Our results suggest that contamination events occurred across the course of collecting and processing the bovine skin and milk samples. Blank controls, including sampling blanks, air blanks and extraction blanks, generated 16S gene copy numbers that were similar to milk samples, highlighting the extremely low microbial biomass of milk from clinically healthy lactating cows and the subsequent challenge in differentiating true from contaminating DNA.

Milk samples (cisternal and stripped milk) generated very low DNA concentration in this study (Figure 2B), which are consistent with general characterization of milk as having a low bacterial biomass with previous reports from cow milk [47,48]. This low DNA content is due in part to several anatomical, physiological, and immunological factors that can help minimize microbial presence within the gland; although microbes can invade the gland and cows do experience mastitis. These factors can also contribute to the presence of large amounts of host somatic cells in milk [22], which can make it challenging to separate DNA from host and microbial sources. Milk is also a technically challenging matrix for microbiome work because its complex chemical composition includes DNase and RNase and other components that can hamper PCR amplification and nucleic acid extraction [14,16,49].

In microbiome research, low biomass samples are especially sensitive to reagent contamination [18,32], which helps explaining the negative correlation we observed between sample biomass (i.e., 16S qPCR copy number) and abundance of contaminants identified by *decontam* (Figure 3E). Our results from both decontam and SourceTracker suggest that contamination of cisternal milk samples mainly occurred during the extraction process (Figures 3C and 4A); and support the presence of bacterial DNA within the extraction kit and PCR reagents. This has been described as “kitome” effect [29,50,51]. Similar conclusions about the impact of extraction have been reported for human milk [28].

The sources of contaminants for stripped milk were more complex than cisternal milk, and included extraction blanks, teat apex, teat canal and cisternal milk samples (Figure 4B). This result is consistent with previous studies [33,52], which suggest that teat apex skin is a source of the microbial population in cow milk. Our results expand on the potential sources of the cow milk microbiome by including technical blanks, air and sampling blanks. During milk sampling and processing, epithelial bacteria can become dislodged and enter the milk being sampled, microbes in the air of the farm or the lab can contaminate the sampling device and/or the milk sample itself, and molecular reagents can contaminate the sample during processing. We note here that the transfer of epithelial and milk-borne bacteria within the teat canal or at the teat apex could be a bidirectional process, particularly if milk leakage is occurring or cows were recently milked. In our sampling scenario, it is impossible to determine the directionality of bacterial transfer events. By defining the skin samples as the source and milk samples as the sink, we are forcing directionality into the analysis and assuming that most of the bacterial transfer is from the skin to the milk. This assumption is partially supported by the fact that cisternal milk did not have the same SourceTracker profile as stripped milk, suggesting that cisternal milk is contaminated as it passes down through the teat canal and out of the teat apex. Sources of this contamination include bacteria within the teat canal, at the teat apex and in the milking parlor environment. To definitively answer this question, additional studies are needed.

It is important to note the very high sample-to-sample variability in the contamination source profile identified by SourceTracker (Figure 4A-B). This high amount of inter-sample variability is typical for low biomass samples and demonstrates the difficulty in measuring the true milk microbiome of lactating dairy cows. We hypothesize that the large variation in the source(s) of milk microbes may be due to the stochastic nature of milk sampling and processing. During these activities, presumably sterile milk can be contaminated with bacteria from the numerous sources we have described in this work. The inherently low bacterial biomass in the milk exacerbates the impact of these events because the bacterial cells involved in a single contamination event can easily overwhelm the number of cells present in the original milk sample. Thus, these mixing and contamination events can easily increase sample-to-sample variability in the profile of source contamination. Library controls were expected to contain very low biomass, and indeed contributed little to the profile of milk and skin sample microbiomes. This only indicates samples are less likely to be contaminated during library preparation but that much of the source of “kitome” contamination occurs prior to library preparation.

### Characterization of the de-contaminated milk microbiome

Even after removal of potential contaminants using both decontam and SourceTracker, the milk samples continued to harbor microbial DNA. Indeed, 53 ASVs remained in cisternal milk and 68 ASVs in stripped milk, though their alpha diversity indexes were lower than those from the skin samples. Stripped milk had similar microbial composition as cisternal milk, which agrees with a previous study that found no significant differences in the microbiota of stripped milk and cisternal milk [24]. Given that stripped milk should represent milk that was removed from the cistern at the time of sampling, we would expect the profiles of cisternal and stripped milk to be similar, particularly after contaminants from the skin and milk parlor air are removed. However, our finding could also stem from a type II error due to our relatively low sample size (N=13 to 14 per sample type) and large variation observed between individual animals. Larger studies will be required to confirm whether cisternal milk and stripped milk microbiota are truly indistinguishable from one another after statistical decontamination.

The profile of the decontaminated milk microbiome data in this study showed that *Staphylococcus* and *Acinetobacter* were the predominant taxa, which is in line with previous studies [53–55]. However, *Bacteroidetes*, *Corynebacterium*, *Pseudomonas* and *Staphylococcus* might be overestimated in previous bovine milk microbiome studies [4,17,24,25,53,56], as they were identified as contaminants in our study. This highlights the importance of contamination removal for correct interpretation of milk microbiome data.

It is also important to note that the 16S sequencing data we analyzed represents DNA from both viable and nonviable bacterial cells. We attempted to discriminate between live and dead bacteria within the milk microbiome data by PMA incorporation into DNA from non-variable bacteria, but the dataset from the PMA-treated samples contained no classifiable bacterial sequence reads. This could indicate that much of the bacterial DNA in the milk microbiome data originated from nonviable bacterial cells, but our culture-based results indicated that 8 of the 28 milk samples did indeed contain viable *Staphylococcus* and *Corynebacterium* bacteria (Table 1). This inability to detect the presence of these bacteria in the PMA data support previous reports that the PMA assay might not be applicable for milk samples [16].

### Potential sources of contamination in bovine teat skin microbiome data

The skin samples (i.e., teat canal and teat apex) exhibited distinct patterns of contamination compared to milk samples, as evidenced by decontam and Sourcetracker (Figure 5A). Results of SourceTracker revealed that the skin samples were mainly contaminated by microbes associated with the sampling devices (as represented by air and sampling blanks, Figure 4A). These blanks were meant to represent all the environmental (i.e., airborne) and physical contact events that sampling devices are exposed to during sample collection and processing. For air blanks (which were opened and waved in the air near the cows), this would include the air in the milking parlor and in the laboratory and biosafety cabinet during sample processing. It would also include physical handling of the device during both sample collection and processing. For the sampling blanks (which were brought to the farm but not opened), these exposure events consisted of the air in the laboratory and biosafety cabinet and handling of the device during sample processing (but not during sample collection). Our study demonstrated that these air and sampling blanks contributed more than 50% of the reads that were generated from teat apex and teat canal samples (Figure 4A), highlighting the importance of aseptic techniques and the inclusion of negative controls during sample collection and processing.

It is important to note here that the environmental microbiome on a dairy farm likely provides a mix of detrimental and beneficial microbes, which could be a biologically important source of udder epithelial microbes. Thus, environmental microbiome is not a source of “contaminants” per se but rather a source of microbes that adhere to and then inhabit the teat apex and/or canal. Indeed, the environmental microbes within dairy farms have been shown to play an important role in the microbial colonization of the udder [57,58]. Therefore, the use of the word “contamination” to describe the sampling device-associated sequences is not strictly precise, because these sequences could represent microbes that were sourced from the barn air and play an important biological or ecological role within the udder microbiome [57]. Further study is needed to discriminate between airborne microbes that contaminate the sample during the actual sampling event, and airborne microbes that actually colonize the epithelium of the mammary gland.

### Statistical approaches for identifying the potential source of sequence reads in bovine teat skin and milk microbiome datasets

We applied two-step statistical algorithms for identification of potential contaminant and source sequences in cow teat epithelium and milk samples, using decontam followed by SourceTracker. Decontam identifies contaminants by considering both the frequency of sequence features and the concentration of sample DNA, as well as the prevalence of sequence features in real samples compared to negative controls [40]. Hence, decontam requires no knowledge of the source of potential contaminants. Instead, the availability of DNA quantification data and proper preparation of negative controls are needed for accurate prediction by decontam. We demonstrated that decontam was robust in detecting contaminants especially in low biomass samples, such as cisternal milk and stripped milk, which accounted for 36% and 27%, respectively, of contaminants in the raw sequences. Furthermore, differences in DNA concentration were correlated with differences in the profile of contaminants (Figure 3), suggesting that sample biomass is a key factor driving the profile of potential contamination [40]. However, contamination is supposed to happen equally in all samples during sample processing, while low biomass samples are at increased risk of contamination because of a lack of competing biological material. The predominant contaminants that we identified using *decontam* include many that have been previously reported as contaminants, i.e., *Pseudomonas*, *Streptococcus*, *Clostridium* and *Turicibacter* [50].

In contrast to decontam, SourceTracker relies on the user to specify both the source and sink samples based on subject-matter knowledge of the study design and sample set, with well-defined contaminating sources providing greater accuracy of prediction than poorly defined source environments [32]. Interestingly, the use of SourceTracker after decontam allowed us to identify many more potential contaminants, suggesting that the approaches used by SourceTracker and decontam are sufficiently distinct to detect different types of contaminating and/or source environment microbes. Indeed, the stated goals of the two methods are distinct and our results suggest that using the two tools in sequence is a robust method to profile the sources of contaminating microbes within low biomass samples such as milk.

It should be noted that SourceTracker assumes that the sink samples are composed wholly of microbes from defined sources. In other words, the sink samples are assumed to be devoid of their own inherent microbiome. As a result, SourceTracker characterizes sequence reads in the sink samples that can’t be associated with a defined source as originating from an “unknown” source. In our analysis, we presumed that these unknown sequences represented the “true” or inherent microbial community within the sink samples (i.e., the milk or skin samples) and therefore we used only these “unknown” sequences for downstream analysis of the true udder and milk microbiomes. While our extensive use of negative controls was comprehensive in terms of capturing most potential sources of contamination, there could be additional sources that we did not capture in our study design, including host sources such as blood and the gastrointestinal tract. These potential sources of milk borne bacteria could be particularly important if the cisternal milk microbiome is being colonized by endogenous transfer from the intestine through the microbiota-gut-mammary axis, especially when the integrity of the milk-blood barrier [59] and the intestinal barrier are impaired by inflammation associated with mastitis [6,60,61]. Furthermore, the negative control samples that we used in this study may not have captured all the bacteria that could have contaminated the milk and skin samples. Additional studies are needed to determine the number and types of negative controls that are needed for sufficiently decontamination of bovine milk microbiome data. It is also important to note that neither decontam nor SourceTracker is intended to identify cross-contamination (i.e., sample-to-sample contamination) that may occur during sampling or through well-to-well leakage during library preparation [62]. A more extensive study of well placement patterns (including many more blank wells) would be needed to identify these cross-contamination events.

### Dynamics of the udder microbiota ecosystem

Similar to previous studies [63,64], teat skin microbiome displayed a more diverse and less dispersed structure as compared to milk samples in this study. That might be due to the exposure of teat skin to a relatively stable biotic or abiotic environments, including air, bedding material, milking system, and milker’s contact [4]. The teat apex is completely exposed to environmental bacteria, while it serves as an incomplete physical barrier against the entry of environmental microbes into the teat canal [65]. We found no significant difference in microbial composition between the teat apex and teat canal samples, supporting the incomplete nature of this physical barrier [66] and indicating that the teat skin microbiome can colonize the teat canal and potentially contaminate the cisternal milk.

### Presence of potential mastitis pathogens with high abundance

After decontamination, we were still able to detect 53 and 68 ASVs representing diverse bacteria in cisternal milk and stripped milk, respectively. However, culture of these milk samples only recovered *Corynebacterium* and *Staphylococcus*, which indicates that the majority of what we detected in the 16S microbiome data was not culturable. This could be because the detected DNA was from nonviable bacteria or because the bacteria either could not grow within our culture conditions or because the bacterial load was below our detection limit of this study (i.e., 100 CFU/mL). The two milk samples grew *Staphylococcus chromogenes* in our culture conditions were from the same cows that had a high relative abundance of *Staphylococcus* in the sequencing data (Figure 8), which indicates that the sequence data does correlate with culture results for culturable bacteria.

We detected an average relative abundance of 14.8% for sequences from common potential mastitis pathogens within our skin and milk samples. Bacterial genera from *Staphylococcus*, *Streptococcus* and *Pseudomonas* were also widely detected in previous studies on teat skin and cow milk microbiome [4,15,67,68]. It is difficult to interpret the relevance of these findings for udder health and dairy cow management because most pathogenic phenotypes are specific to certain species or even strains of bacteria while traditional 16S approaches we and others have used are only able to classify most sequences to the family or genus levels. However, given the importance of these potential mastitis pathogens in the development of intramammary infections and the importance of understanding whether and/or how microbiome results can be associated with udder health, additional more in-depth studies of this nature are warranted.

## Conclusions

After quality control and removal of potential contaminants identified by decontam and SourceTracker, only 20-24% of the identified sequence reads remained for udder skin and milk microbiome analysis in this study. Source of contamination differed among sample types. Specifically, milk samples were mainly contaminated by DNA from the extraction kit and skin while teat skin microbiomes were mainly contaminated from microorganisms in the air and from the sampling devices. No significant differences in microbial profiles were detected in decontaminated milk samples (cisternal milk vs. stripped milk) or in decontaminated skin samples (teat apex vs. teat canal). However, milk samples displayed a less diverse, more dispersed, and compositionally distinct microbial profile compared to teat skin samples. Across all milk and skin samples, *Actinobacteria* and *Staphylococcus* were the predominant genera and on average 14.8% of the sequence reads were classified as being from potential mastitis pathogens. In contract, culture of milk samples only detected the presence of *Staphylococcus* in one cisternal milk sample and *Corynebacterium* and *Staphylococcus* in 50% of stripped milk samples. Our PMA-treated milk samples did not yield any classable sequence reads except for host DNA. Our results highlight the importance of aseptic sampling for udder skin microbiome studies. For future research, we strongly recommend the inclusion of sufficient negative controls to identify potential contamination events and to generate more reliable microbiome data.

## Declaration

### Ethics approval and Consent to participate

All the animal procedures and activities involved in this study were approved by the University of Minnesota Institutional Animal Care and Use Committee (protocol number 1904- 36973A).

### Consent for publication

Not applicable.

### Availability of data and materials

The sequencing data of this article are available at National Center for Biotechnology Information (NCBI) Sequence Read Archive (SRA) under the Bioproject accession number PRJNA983382 (https://www.ncbi.nlm.nih.gov/bioproject/?term=PRJNA983382). The metadata and R scripts necessary to reproduce the analysis presented in this paper are available at https://github.com/TheNoyesLab/Bovine_Milk_Microbiome.

### Authors’ contributions

CJD, LSC and NRN conceived of the project and obtained funding for the project. BC oversaw the animals sampled as part of the project. CJD, TCW, FPM and LSC collected samples for the project. TCW processed samples for PMA analysis with oversight by TR. CJD and TR processed samples for microbiome analysis. CJD processed samples for microbiological culture submission. CJD and YD performed bioinformatic analysis with input from NRN. YD performed statistical analysis and interpretation with input from NRN. YD wrote the main manuscript. YD and CJD prepared figures. All authors provided input on the manuscript and interpretation of results.

## Acknowledgements

The authors acknowledge the service provided by Udder Health Lab at the University of Minnesota for milk culture results and the University of Minnesota Dairy Cattle Teaching and Research Center for the support with the animal portion of this trial and cow management. We would like to thank the University of Minnesota Genomics Center for amplicon sequencing and the Minnesota Supercomputing Institute (MSI) at the University of Minnesota for sequencing data storage.

## References

1. Mariadassou M, Nouvel LX, Constant F, Morgavi DP, Rault L, Barbey S, et al. Microbiota members from body sites of dairy cows are largely shared within individual hosts throughout lactation but sharing is limited in the herd. Animal Microbiome. 2023;5:32.

2. Hanning I, Diaz-Sanchez S. The functionality of the gastrointestinal microbiome in non-human animals. Microbiome. 2015;3:51.

3. Hooper LV, Littman DR, Macpherson AJ. Interactions Between the Microbiota and the Immune System. Science. 2012;336:1268–73.

4. Derakhshani H, Fehr KB, Sepehri S, Francoz D, De Buck J, Barkema HW, et al. Invited review: Microbiota of the bovine udder: Contributing factors and potential implications for udder health and mastitis susceptibility. Journal of Dairy Science. 2018;101:10605–25.

5. Quigley L, O’Sullivan O, Stanton C, Beresford TP, Ross RP, Fitzgerald GF, et al. The complex microbiota of raw milk. FEMS Microbiology Reviews. 2013;37:664–98.

6. Addis MF, Tanca A, Uzzau S, Oikonomou G, Bicalho RC, Moroni P. The bovine milk microbiota: insights and perspectives from-omics studies. Mol BioSyst. 2016;12:2359–72.

7. Rainard P. Mammary microbiota of dairy ruminants: fact or fiction? Veterinary Research. 2017;48:25.

8. Parente E, Ricciardi A, Zotta T. The microbiota of dairy milk: A review. International Dairy Journal. 2020;107:104714.

9. Taponen S, McGuinness D, Hiitiö H, Simojoki H, Zadoks R, Pyörälä S. Bovine milk microbiome: a more complex issue than expected. Veterinary Research. 2019;50:44.

10. Weinroth MD, Belk AD, Dean C, Noyes N, Dittoe DK, Rothrock MJ Jr, et al. Considerations and best practices in animal science 16S ribosomal RNA gene sequencing microbiome studies. Journal of Animal Science. 2022;100:skab346.

11. Rault L, Lévêque P-A, Barbey S, Launay F, Larroque H, Le Loir Y, et al. Bovine Teat Cistern Microbiota Composition and Richness Are Associated With the Immune and Microbial Responses During Transition to Once-Daily Milking. Frontiers in Microbiology. 2020;11. Available from: https://www.frontiersin.org/articles/10.3389/fmicb.2020.602404

12. Wouters JTM, Ayad EHE, Hugenholtz J, Smit G. Microbes from raw milk for fermented dairy products. International Dairy Journal. 2002;12:91–109.

13. Guo W, Liu S, Khan MZ, Wang J, Chen T, Alugongo GM, et al. Bovine milk microbiota: Key players, origins, and potential contributions to early-life gut development. Journal of Advanced Research. 2023; Available from: https://www.sciencedirect.com/science/article/pii/S2090123223001790

14. Foroutan A, Guo AC, Vazquez-Fresno R, Lipfert M, Zhang L, Zheng J, et al. Chemical Composition of Commercial Cow’s Milk. J Agric Food Chem. 2019;67:4897–914.

15. Dean CJ, Slizovskiy IB, Crone KK, Pfennig AX, Heins BJ, Caixeta LS, et al. Investigating the cow skin and teat canal microbiomes of the bovine udder using different sampling and sequencing approaches. Journal of Dairy Science. 2021;104:644–61.

16. Ganda E, Beck KL, Haiminen N, Silverman JD, Kawas B, Cronk BD, et al. DNA Extraction and Host Depletion Methods Significantly Impact and Potentially Bias Bacterial Detection in a Biological Fluid. mSystems. 2021;6:e00619–21.

17. Dahlberg J, Sun L, Waller KP, Östensson K, McGuire M, Agenäs S, et al. Microbiota data from low biomass milk samples is markedly affected by laboratory and reagent contamination. PLOS ONE. 2019;14:e0218257.

18. Eisenhofer R, Minich JJ, Marotz C, Cooper A, Knight R, Weyrich LS. Contamination in Low Microbial Biomass Microbiome Studies: Issues and Recommendations. Trends in Microbiology. 2019;27:105–17.

19. Pollock J, Salter SJ, Nixon R, Hutchings MR. Milk microbiome in dairy cattle and the challenges of low microbial biomass and exogenous contamination. anim microbiome. 2021;3:80.

20. Weiss S, Amir A, Hyde ER, Metcalf JL, Song SJ, Knight R. Tracking down the sources of experimental contamination in microbiome studies. Genome Biology. 2014;15:564.

21. Weyrich LS, Farrer AG, Eisenhofer R, Arriola LA, Young J, Selway CA, et al. Laboratory contamination over time during low-biomass sample analysis. Molecular Ecology Resources. 2019;19:982–96.

22. Dean CJ, Peña-Mosca F, Ray T, Heins BJ, Machado VS, Pinedo PJ, et al. Evaluation of Contamination in Milk Samples Pooled From Independently Collected Quarters Within a Laboratory Setting. Frontiers in Veterinary Science. 2022;9. Available from: https://www.frontiersin.org/articles/10.3389/fvets.2022.818778

23. Adkins PR, Middleton JR. Laboratory handbook on bovine mastitis. National Mastitis Council, Incorporated; 2017.

24. Dahlberg J, Williams JE, McGuire MA, Peterson HK, Östensson K, Agenäs S, et al. Microbiota of bovine milk, teat skin, and teat canal: Similarity and variation due to sampling technique and milk fraction. Journal of Dairy Science. 2020;103:7322–30.

25. Metzger SA, Hernandez LL, Skarlupka JH, Walker TM, Suen G, Ruegg PL. A Cohort Study of the Milk Microbiota of Healthy and Inflamed Bovine Mammary Glands From Dryoff Through 150 Days in Milk. Frontiers in Veterinary Science. 2018;5. Available from: https://www.frontiersin.org/articles/10.3389/fvets.2018.00247

26. Neave F. Diagnosis of mastitis by bacteriological methods alone. Doc Int Dairy Fed. 1975;

27. Jervis-Bardy J, Leong LEX, Marri S, Smith RJ, Choo JM, Smith-Vaughan HC, et al. Deriving accurate microbiota profiles from human samples with low bacterial content through post-sequencing processing of Illumina MiSeq data. Microbiome. 2015;3:19.

28. Moossavi S, Fehr K, Khafipour E, Azad MB. Repeatability and reproducibility assessment in a large-scale population-based microbiota study: case study on human milk microbiota. Microbiome. 2021;9:41.

29. Salter SJ, Cox MJ, Turek EM, Calus ST, Cookson WO, Moffatt MF, et al. Reagent and laboratory contamination can critically impact sequence-based microbiome analyses. BMC Biology. 2014;12:87.

30. Hornung BVH, Zwittink RD, Kuijper EJ. Issues and current standards of controls in microbiome research. FEMS Microbiology Ecology. 2019;95:fiz045.

31. Zaramela LS, Tjuanta M, Moyne O, Neal M, Zengler K. synDNA—a Synthetic DNA Spike-in Method for Absolute Quantification of Shotgun Metagenomic Sequencing. mSystems. 2022;7:e00447–22.

32. Karstens L, Asquith M, Davin S, Fair D, Gregory WT, Wolfe AJ, et al. Controlling for Contaminants in Low-Biomass 16S rRNA Gene Sequencing Experiments. mSystems. 2019;4:e00290–19.

33. Doyle CJ, Gleeson D, O’Toole PW, Cotter PD. Impacts of Seasonal Housing and Teat Preparation on Raw Milk Microbiota: a High-Throughput Sequencing Study. Applied and Environmental Microbiology. 2016;83:e02694–16.

34. Rowe SM, Godden SM, Royster E, Timmerman J, Boyle M. Cross-sectional study of the relationship between cloth udder towel management, towel bacteria counts, and intramammary infection in late-lactation dairy cows. Journal of Dairy Science. 2019;102:11401–13.

35. Jahan NA, Godden SM, Royster E, Schoenfuss TC, Gebhart C, Timmerman J, et al. Evaluation of the matrix-assisted laser desorption ionization time of flight mass spectrometry (MALDI-TOF MS) system in the detection of mastitis pathogens from bovine milk samples. Journal of Microbiological Methods. 2021;182:106168.

36. Wickham H. Data Analysis. In: Wickham H, editor. ggplot2: Elegant Graphics for Data Analysis. Cham: Springer International Publishing; 2016. p. 189–201. Available from: 10.1007/978-3-319-24277-4_9

37. Callahan BJ, McMurdie PJ, Rosen MJ, Han AW, Johnson AJA, Holmes SP. DADA2: High-resolution sample inference from Illumina amplicon data. Nat Methods. 2016;13:581–3.

38. McMurdie PJ, Holmes S. phyloseq: An R Package for Reproducible Interactive Analysis and Graphics of Microbiome Census Data. PLOS ONE. 2013;8:e61217.

39. Altschul SF, Gish W, Miller W, Myers EW, Lipman DJ. Basic local alignment search tool. Journal of Molecular Biology. 1990;215:403–10.

40. Davis NM, Proctor DM, Holmes SP, Relman DA, Callahan BJ. Simple statistical identification and removal of contaminant sequences in marker-gene and metagenomics data. Microbiome. 2018;6:226.

41. Knights D, Kuczynski J, Charlson ES, Zaneveld J, Mozer MC, Collman RG, et al. Bayesian community-wide culture-independent microbial source tracking. Nat Methods. 2011;8:761–3.

42. Wisconsin Veterinary Diagnostic Laboratory. Interpretation of mastitis milk culture results. 2016. Available from: https://www.wvdl.wisc.edu/wp-content/uploads/2016/07/Interpretation-of-Mastitis-Culture-Results-16-07-15.pdf

43. Tiedemann F. gghalves: Compose Half-Half Plots Using Your Favourite Geoms. 2022. Available from: https://cran.r-project.org/web/packages/gghalves/index.html

44. Anderson MJ. A new method for non-parametric multivariate analysis of variance. Austral Ecology. 2001;26:32–46.

45. Oksanen J, Simpson GL, Blanchet FG, Kindt R, Legendre P, Minchin PR, et al. vegan: Community Ecology Package. 2022. Available from: https://cran.r-project.org/web/packages/vegan/index.html

46. Lenth RV, Bolker B, Buerkner P, Giné-Vázquez I, Herve M, Jung M, et al. emmeans: Estimated Marginal Means, aka Least-Squares Means. 2023. Available from: https://cran.r-project.org/web/packages/emmeans/index.html

47. Sun L, Dicksved J, Priyashantha H, Lundh Å, Johansson M. Distribution of bacteria between different milk fractions, investigated using culture-dependent methods and molecular-based and fluorescent microscopy approaches. Journal of Applied Microbiology. 2019;127:1028–37.

48. Schwenker JA, Friedrichsen M, Waschina S, Bang C, Franke A, Mayer R, et al. Bovine milk microbiota: Evaluation of different DNA extraction protocols for challenging samples. MicrobiologyOpen. 2022;11:e1275.

49. Soboleva SE, Zakharova OD, Sedykh SE, Ivanisenko NV, Buneva VN, Nevinsky GA. DNase and RNase activities of fresh cow milk lactoferrin. Journal of Molecular Recognition. 2019;32:e2777.

50. Glassing A, Dowd SE, Galandiuk S, Davis B, Chiodini RJ. Inherent bacterial DNA contamination of extraction and sequencing reagents may affect interpretation of microbiota in low bacterial biomass samples. Gut Pathogens. 2016;8:24.

51. Kim D, Hofstaedter CE, Zhao C, Mattei L, Tanes C, Clarke E, et al. Optimizing methods and dodging pitfalls in microbiome research. Microbiome. 2017;5:52.

52. Verdier-Metz I, Gagne G, Bornes S, Monsallier F, Veisseire P, Delbès-Paus C, et al. Cow Teat Skin, a Potential Source of Diverse Microbial Populations for Cheese Production. Applied and Environmental Microbiology. 2012;78:326–33.

53. Breitenwieser F, Doll EV, Clavel T, Scherer S, Wenning M. Complementary Use of Cultivation and High-Throughput Amplicon Sequencing Reveals High Biodiversity Within Raw Milk Microbiota. Frontiers in Microbiology. 2020;11. Available from: https://www.frontiersin.org/articles/10.3389/fmicb.2020.01557

54. Cremonesi P, Severgnini M, Romanò A, Sala L, Luini M, Castiglioni B. Bovine Milk Microbiota: Comparison among Three Different DNA Extraction Protocols To Identify a Better Approach for Bacterial Analysis. Microbiology Spectrum. 2021;9:e00374–21.

55. Kable ME, Srisengfa Y, Laird M, Zaragoza J, McLeod J, Heidenreich J, et al. The Core and Seasonal Microbiota of Raw Bovine Milk in Tanker Trucks and the Impact of Transfer to a Milk Processing Facility. mBio. 2016;7:e00836–16.

56. Lima SF, Teixeira AGV, Lima FS, Ganda EK, Higgins CH, Oikonomou G, et al. The bovine colostrum microbiome and its association with clinical mastitis. Journal of Dairy Science. 2017;100:3031–42.

57. Hohmann M-F, Wente N, Zhang Y, Krömker V. Bacterial Load of the Teat Apex Skin and Associated Factors at Herd Level. Animals. 2020;10:1647.

58. Rowbotham RF, Ruegg PL. Bacterial counts on teat skin and in new sand, recycled sand, and recycled manure solids used as bedding in freestalls. Journal of Dairy Science. 2016; 99(8):6594–6608.

59. Wellnitz O, Bruckmaier RM. Invited review: The role of the blood-milk barrier and its manipulation for the efficacy of the mammary immune response and milk production. J Dairy Sci. 2021;104:6376–88.

60. Young W, Hine BC, Wallace OAM, Callaghan M, Bibiloni R. Transfer of intestinal bacterial components to mammary secretions in the cow. PeerJ. 2015;3:e888.

61. Zhao C, Hu X, Qiu M, Bao L, Wu K, Meng X, et al. Sialic acid exacerbates gut dysbiosis-associated mastitis through the microbiota-gut-mammary axis by fueling gut microbiota disruption. Microbiome. 2023;11:78.

62. Austin GI, Park H, Meydan Y, Seeram D, Sezin T, Lou YC, et al. Contamination source modeling with SCRuB improves cancer phenotype prediction from microbiome data. Nat Biotechnol. 2023;1–9.

63. Andrews T, Neher DA, Weicht TR, Barlow JW. Mammary microbiome of lactating organic dairy cows varies by time, tissue site, and infection status. PLOS ONE. 2019;14:e0225001.

64. Derakhshani H, Plaizier JC, De Buck J, Barkema HW, Khafipour E. Composition and co-occurrence patterns of the microbiota of different niches of the bovine mammary gland: potential associations with mastitis susceptibility, udder inflammation, and teat-end hyperkeratosis. Animal Microbiome. 2020;2:11.

65. Hogan J, Smith KL. Managing environmental mastitis. Vet Clin North Am Food Anim Pract. 2012;28(2):217–224.

66. Zecconi A, Hamanno J, Bronzo V, Moroni P, Giovannini G, Piccinini R. Relationship Between Teat Tissue Immune Defences and Intramammary Infections. In: Mol JA, Clegg RA, editors. Biology of the Mammary Gland. Boston, MA: Springer US; 2002. p. 287–93. Available from: 10.1007/0-306-46832-8_33

67. Hiitiö H, Simojoki H, Kalmus P, Holopainen J, Pyörälä S, Taponen S. The effect of sampling technique on PCR-based bacteriological results of bovine milk samples. Journal of Dairy Science. 2016;99:6532–41.

68. Frétin M, Martin B, Rifa E, Isabelle V-M, Pomiès D, Ferlay A, et al. Bacterial community assembly from cow teat skin to ripened cheeses is influenced by grazing systems. Sci Rep. 2018;8:200.

